# TRIDENT (Taxonomic Resolution and IDentification using Environmental dNa Traces): An Optimized Algorithm for Vertebrate Taxonomic Assignments in eDNA Metabarcoding, Integrating Molecular, Taxonomic, and Ecological Criteria

**DOI:** 10.64898/2026.06.29.735257

**Authors:** Rachel Haderlé, Gabriel Jung, Manon Riou, Visotheary Ung, Jean-Luc Jung

## Abstract

Environmental DNA (eDNA) metabarcoding has become a powerful approach for large-scale biodiversity assessment, yet taxonomic assignment remains one of its most critical error-prone steps. Current bioinformatic pipelines rely on molecular similarity searches against reference databases, but assignment accuracy is constrained not only by short marker length and database incompleteness, but also by fundamental limitations, including recent species radiations, incomplete lineage sorting, introgression, NUMTs, and the imperfect correspondence between genetic variation and species boundaries.

Here, we present TRIDENT (Taxonomic Resolution and IDentification using Environmental dNa Traces), an automated and simple protocol designed to improve taxonomic assignments in eDNA metabarcoding. Developed for marine vertebrates, TRIDENT may be used with any barcode and integrates three complementary sources of evidence: molecular similarity (NCBI/GenBank and BOLD), curated taxonomic information (WoRMS), and ecological plausibility derived from biogeographic occurrence data (GBIF). The workflow sequentially constructs candidate taxon lists based on sequence similarity, expands them through taxonomic hierarchies, and filters them using spatial occurrence constraints. It further identifies possible taxa lacking reference barcodes and evaluates their plausibility through CO1-based similarity if data exist in BOLD.

TRIDENT has been implemented as a source-available Python tool and tested using empirical eDNA datasets from marine vertebrates as well as simulated communities. Results demonstrate that the tool produces taxonomic assignments consistent with expert manual curation while substantially reducing processing time and attention errors caused by manual processing of large datasets. By combining molecular, taxonomic, and ecological criteria within a single framework, TRIDENT improves transparency and reproducibility and provides a robust and flexible solution strengthening confidence in taxonomic identifications in eDNA-based biodiversity assessments.

## INTRODUCTION

Molecular barcoding has profoundly improved species identification by providing a DNA-based framework for taxonomic assignment. DNA barcoding *sensu stricto* was initially established in animals using a standardized genetic marker, referred to as a “barcode”, corresponding to approximately 500bp of the 5’ end of the mitochondrial cytochrome c oxidase subunit 1 (CO1) gene (Hebert et al. 2003). This approach enables rapid, standardized, and broadly applicable species-level identification across diverse taxa (Brown et al. 1999; Bucklin et al. 1999; Vincent et al. 2000; Trewick 2000; Alfonsi et al. 2013; Jung 2017). DNA barcoding *sensu lato* refers more broadly to the identification of any taxonomic rank using the sequence of any DNA fragment, without restriction to a specific marker or taxonomic resolution (Valentini et al. 2009). Overall, the DNA barcoding concept was developed to accelerate biodiversity assessments and inventories and to support conservation and management actions (Schander and Willassen 2005; Ball and Armstrong 2006; Neigel et al. 2007; Hajibabaei et al. 2007a; Stoeckle and Hebert 2008; Swartz et al. 2008; Krishna Krishnamurthy and Francis 2012; Jung et al. 2016).

Advances in high-throughput sequencing technologies have enabled the development of DNA metabarcoding, which allows the simultaneous detection of multiple taxa from bulk biological samples or environmental DNA (eDNA) extracts (Taberlet et al. 2012). By targeting short DNA fragments as barcodes and sequencing millions of reads in parallel, metabarcoding has opened new avenues for large-scale biodiversity surveys. However, despite its transformative potential, taxonomic assignment remains one of the most critical and error-prone steps in metabarcoding workflows (Calderón-Sanou et al. 2020; Blackman et al. 2023; Keck et al. 2023; Watts et al. 2026).

Molecular taxonomic assignment methods can be broadly classified into four categories: similarity-based, composition-based, phylogeny-based, and probabilistic approaches, whose performance varies widely depending on methodological and biological contexts (Bazinet and Cummings 2012; Pitz et al. 2020; Hleap et al. 2021). These differences stem from both conceptual and technical limitations inherent to molecular identification. Notably, taxonomic units are generally not defined on the basis of DNA sequences, making the correspondence between molecular variation and taxonomic boundaries inherently imperfect (Miralles et al. 2024). This challenge is further exacerbated in metabarcoding studies, which typically rely on short barcodes (ca. 60–250 bp) that often lack sufficient phylogenetic signal for fine taxonomic resolution.

As a result, molecular taxonomic assignment accuracy is strongly shaped by multiple interacting factors, including sample processing, primer selection, PCR conditions, barcode choice, and the classification algorithm used (Bazinet and Cummings 2012; Aylagas et al. 2016; Clarke et al. 2017; Lobo et al. 2017; Pitz et al. 2020). In particular, the different genetic markers used as barcodes may mutate at different evolutionary rates and exhibit distinct evolutionary dynamics, leading to variable discriminatory power among taxa (Suárez-Menéndez et al. 2023). A central assumption of DNA barcoding is the existence of a “barcoding gap”, defined as a clear separation between intra-and interspecific genetic distances (Hebert et al. 2004; Wiemers and Fiedler 2007). In principle, this separation allows sequences to be assigned to species based on their relative genetic similarity to reference sequences. However, the existence of a distinct barcoding gap is not universal and may be compromised when using short metabarcoding fragments, thereby reducing assignment reliability (Hajibabaei et al. 2007b; Alfonsi et al. 2013; Claver et al. 2023). Moreover, biological and/or evolutionary processes, such as recent species divergence, hybridization (Amaral et al. 2014), introgression (Lehman et al. 1991; Verginelli et al. 2005), high intraspecific variability (Meier et al. 2006), heteroplasmy (Vollmer et al. 2011) and NUMT (Nuclear Mitochondrial DNA; Ko et al. 2015) can further obscure taxonomic boundaries (Vinas and Tudela 2009; Alfonsi et al. 2013).

Beyond marker-related constraints, taxonomic assignments are highly dependent on reference databases, which are often incomplete, geographically biased, or affected by taxonomic misidentifications (Hleap et al. 2021; Meglécz 2023). While curated, local, or ecologically filtered reference databases can substantially improve assignment accuracy (Gold et al. 2021; Blackman et al. 2023; Meglécz 2023), their limited taxonomic coverage may hinder the detection of invasive, cryptic, or range-expanding species. Consequently, the integration of expert taxonomic knowledge, ecological information, and regional species inventories remains essential for ensuring robust and biologically meaningful interpretations of metabarcoding data (Coissac et al. 2012; Deiner et al. 2017; Blackman et al. 2023).

Environmental DNA (eDNA) metabarcoding, which couples the analysis of DNA traces shed into the environment with high-throughput sequencing, has emerged as a powerful, non-invasive, and scalable tool for biodiversity assessment (Valentini et al. 2009; Deiner et al. 2017; Taberlet et al. 2018; Afzali et al. 2021; Pochon et al. 2024). This approach enables the detection of rare, elusive, or cryptic species that are often overlooked by conventional survey methods (Takahashi et al. 2023; Bernatchez et al. 2024) and is increasingly adopted in biomonitoring programs worldwide (Deiner et al. 2017; Pawlowski et al. 2020; Jung 2024; Haderlé et al. 2024, 2026a, c; Watts et al. 2026; Madon et al. 2025).

The reliability of eDNA-based inventories, however, strongly depends on transparent, reproducible, and well-documented bioinformatic workflows (Hleap et al. 2021; Keck et al. 2023). In practice, taxonomic assignments generated by automated pipelines are often used directly in downstream analyses, with limited scrutiny of their ecological plausibility or consistency with known species distributions (Kaehler et al. 2019; Blackman et al. 2023). eDNA metabarcoding datasets typically comprise tens to hundreds of thousands of sequences reads per sample, which bioinformatic pipelines denoise and cluster into molecular units, commonly referred to as OTUs, MOTUs, or ASVs (Hakimzadeh et al. 2024; **Appendix 1**). These units are then automatically assigned to taxa and treated as biologically meaningful entities. Yet several sources of bias may compromise the validity of these assignments. Errors or misannotations in reference databases and the co-amplification of NUMTs can lead to inaccurate or inflated diversity estimates (Lopez et al. 2022; Blackman et al. 2023). In addition, the lack of integration of biogeographic information and incomplete reference libraries may result in ecologically implausible or overconfident species-level identifications, particularly when local taxa are absent from databases.

To address limitations of automated taxonomic assignment, we developed TRIDENT (Taxonomic Resolution and IDentification using Environmental dNa Traces). TRIDENT is constructed on an optimized algorithm for marine vertebrate eDNA (M)OTUs/ASVs taxonomic automated assignments, based on three pillars: (*i*) it integrates molecular similarity metrics, (*ii*) it uses curated taxonomic reference lists, and (*iii*) it takes into account ecological plausibility criteria. Importantly, TRIDENT is designed to be gene-agnostic and can be applied to vertebrate metabarcoding datasets regardless of the genetic marker or barcode used. It also identifies inconsistent (M)OTUs/ASVs, such as sequencing artefacts, NUMTs, or exogenous contamination. TRIDENT accounts for taxa lacking reference barcodes, flagging them as potentially valid but unresolved. It can also take advantage of molecular data coming from the presently more exhaustive reference database for animal molecular taxonomy, e.g. BOLD (Ratnasingham and Hebert 2007; Ratnasingham et al. 2024), even if the user-chosen barcode is not CO1. By linking molecular, taxonomic, and ecological evidence in an automated process, and while saving time for those conducting experiments, TRIDENT improves eDNA metabarcoding assignment reliability and strengthens downstream biodiversity analyses.

## MATERIALS & METHODS

### 1. General Architecture and Algorithm of TRIDENT

TRIDENT relies on several recognized external databases and consists of three successive steps (**Fig. 1**). The protocol was initially developed iteratively on an eDNA dataset of marine vertebrate samples collected in Guadeloupe waters (Haderlé et al. 2024).

**Figure 1:**
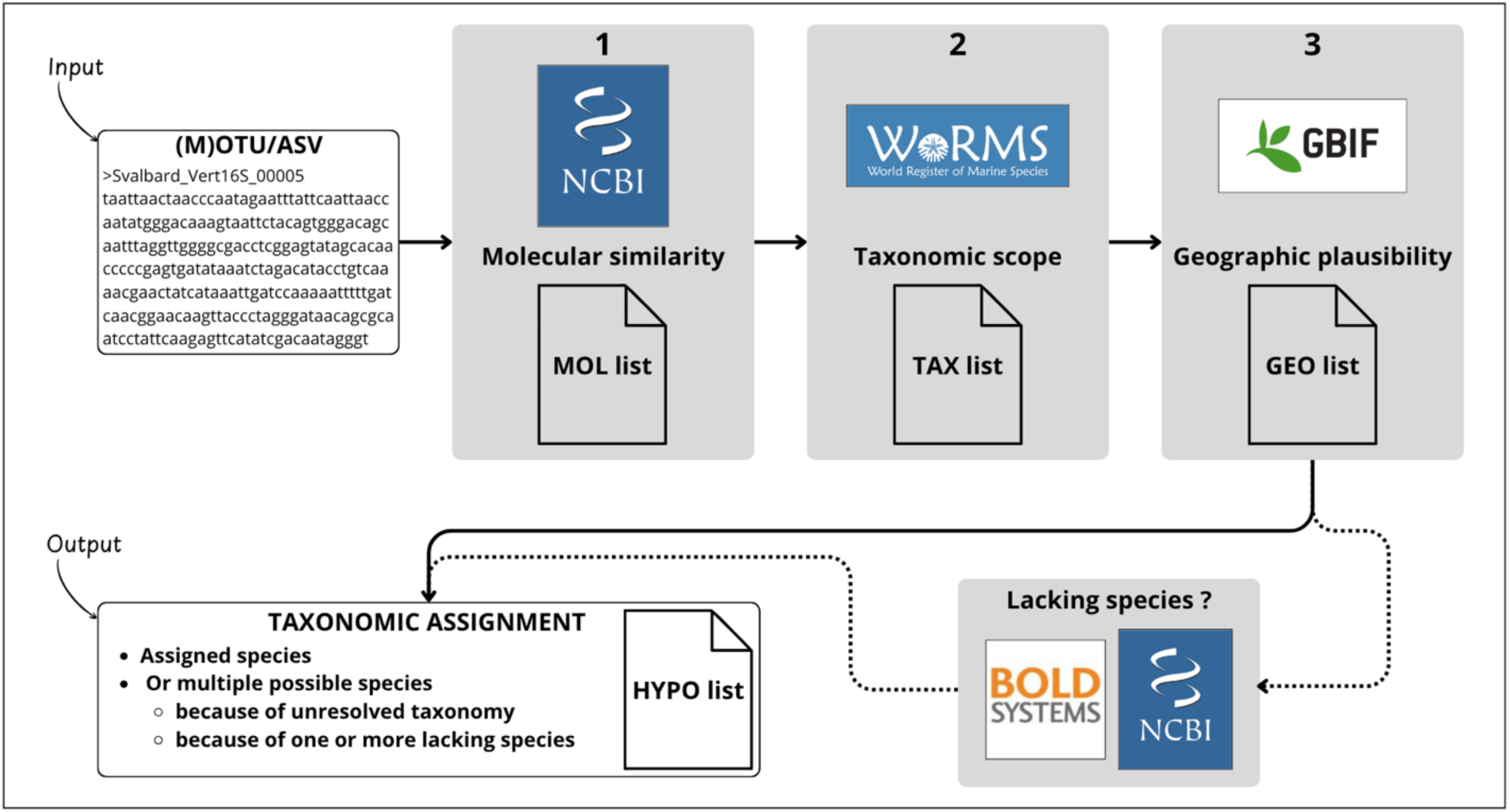
Summary of the main stages of the TRIDENT protocol; white boxes represent protocol inputs and outputs, while grey boxes correspond to individual processing stages, together with the database(s) queried and the output list produced at each stage; solid arrows indicate the core workflow of the protocol and dashed arrows represent optional supplementary steps.

#### a. First step: the MOL list

The first step consists of comparing each (M)OTU/ASV sequence against the NCBI Core Nucleotide Database (GenBank) using BLASTn (Altschul et al. 1990). Users can modify the search parameters. TRIDENT analyses BLAST results to detect a possible barcoding gap with a user-adjustable threshold set by default to 2% of sequence divergence. When a barcoding gap is detected, it is used to delimit the range of candidate taxa. By default, this approach is complemented by a similarity threshold, which retains taxa falling within the top 2% similarity range. Alternatively, users can select the similarity threshold method alone, without applying the barcoding gap approach. Both thresholds are customizable. The resulting list of potential species (MOL list) is accompanied by a graphical representation showing the distribution of similarity values and highlighting the position of the detected barcoding gap, when present.

#### b. Second step: the TAX list

The second step consists of querying the World Register of Marine Species – WoRMS (Costello et al. 2013) to extend the list to include all accepted species belonging to the genera represented by genus- or species-level BLAST hits retained in the MOL list, thereby generating the TAX list. Species excluded during the first step (MOL list) are not retained in the TAX list.

#### c. Third step: the GEO list

The third step consists of evaluating the geographic plausibility of all species in the TAX list using occurrence data from the Global Biodiversity Information Facility – GBIF (GBIF: The Global Biodiversity Information Facility 2026). A geographic search window is defined using user-configured latitude, longitude, and radius, and the area is also represented graphically. Species are retained only if they occur within this area, with a customizable minimum number of occurrences used to reduce erroneous records (default: n = 3). All retained species constitute the GEO list.

### 2. Cases of Species not Represented in GenBank

Species appearing in both TAX and GEO lists but absent from the MOL list constitute a special case and are treated as candidate taxa lacking molecular references. When available, their CO1 barcode sequences are retrieved from the Barcode of Life Data System (BOLD). Among these, the most divergent CO1 sequences are retained in order to maximize intraspecific variability. Similarly, CO1 sequences for species in the MOL list are searched and retained for comparison with previous ones using BLAST.

This step assumes that high CO1 similarities are likely to reflect a comparable similarity for the metabarcoding marker used, whereas low CO1 similarity indicates unlikely species correspondence. Species are retained if CO1 sequence similarity exceeds a user-defined threshold (default: 95% identity), and if the targeted marker is confirmed to be absent from GenBank. Whenever a taxon is confirmed to lack the target barcode in GenBank, yet is present in both the TAX and GEO lists, and shows high CO1 similarity to a closely related taxon retained in the MOL list, it is classified as a hypothetical taxon.

### 3. Output of TRIDENT

TRIDENT generates an overview table summarizing results for all (M)OTUs/ASVs. Each row corresponds to a species assigned to a given (M)OTU/ASV and includes comprehensive metadata documenting every processing step for each taxon, ensuring transparency and traceability. In addition, a Sankey diagram is produced for each (M)OTU/ASV, showing how candidate taxa progress through the successive processing steps. Users can then select the taxa to retain for the final assignment. This produces a second output table with one row per (M)OTU/ASV and the assigned taxon. When assignment at the species level is not possible, all plausible species are reported in the “identificationRemarks” column.

Output fields comply with FAIR standards adapted to eDNA datasets (Abarenkov et al. 2023; Takahashi et al. 2025). The result interface also provides detailed information for each (M)OTU/ASV, as well as for each species listed in the overview table, with direct access to external resources such as WoRMS, GBIF and NCBI.

### 4. Implementation of TRIDENT

TRIDENT is a source-available Python package (available at: https://github.com/ISYEB-ADNe/TRIDENT) under the PolyForm Strict License 1.0.0 running on Linux, macOS and Windows and installable with a single command. It provides a web-based graphical interface that guides the user through the successive steps of TRIDENT (**Fig. 1**), requiring no programming expertise.

The input is a standard FASTA file containing one or several (M)OTU/ASV, and the main output is a formatted list of taxon assignments for each (M)OTU/ASV. At each step, specific summaries are displayed to follow the current state of the analysis and help refine the configurable parameters, and all intermediate results can be exported as *.csv* files. The complete pipeline state ((M)OTU/ASV, parameters and results) is stored in a single SQLite database that can itself be shared and reused as an alternative input, ensuring full reproducibility.

Internally, TRIDENT is organised into three layers: a set of API clients, the pipeline module that orchestrates the protocol, and the user interface. The client layer queries external biodiversity databases and processes the returned data. NCBI is searched with BLASTn through Biopython (Cock et al. 2009) to identify similar sequences and derive a list of candidate species (https://biopython.org/docs/dev/api/Bio.Blast.NCBIWWW.html). The WoRMS REST API (https://www.marinespecies.org/rest/) expands each genus to its accepted marine species. The GBIF REST API (https://techdocs.gbif.org/en/openapi/) counts the occurrences of a given taxon within a user-defined geographical bounding box. Finally, the BOLD REST API (https://boldsystems.org/data/api/) provides CO1 proxy sequences for the indirect validation of candidate species found in GBIF but absent from NCBI. For each of those databases, relevant filters on the raw results have been constructed (e.g. a barcoding gap filter for the BLAST results, and a selector of sufficiently different sequences obtained from BOLD to reduce their number from typically hundreds to about ten per species).

The pipeline layer runs the protocol in sequence: each step uses the output of the previous one and executes both the database search and the filtering of raw results, from the initial BLAST search to the final taxon list. For example, the NCBI BLAST search yields a set of candidate species, from which the corresponding genera are extracted after applying a barcoding gap or similarity filter and serve as input for the WoRMS query. All intermediate results are written to the local database created automatically for the input FASTA file, and a caching system ensures that rerunning a step with modified parameters only re-queries the data that have actually changed, avoiding redundant calls to the external databases (in the example above, if a slightly looser filter on the BLAST hits adds only one or two genera, only those are queried in WoRMS).

The user interface exposes the parameters relevant to each step and presents its results through dedicated metrics, summary tables and figures, making anomalous, incomplete or validated results easy to spot while keeping the raw outputs available for closer inspection. A final panel summarises the validated species list and lets the user manually include or exclude taxa, refining the output beyond the automated filters.

### 5. Performance Testing of TRIDENT

A series of complementary tests were conducted to evaluate the performance of TRIDENT. The first test series was performed on selected MOTUs derived from two environmental DNA (eDNA) metabarcoding surveys of marine vertebrates. The first dataset (Haderlé et al. 2026b) originated from Guadeloupe (Guadeloupe dataset: https://www.gbif.org/dataset/fb6939d1-0b75-497e-a54b-2a38a907bf05), and the second (Haderlé et al. 2026c) from Svalbard (Svalbard dataset: https://www.gbif.org/dataset/8c8f29d8-870b-44fd-82c0-c767734e0f27). For the Guadeloupe dataset, amplification targeted a ∼97 bp fragment of the mitochondrial 12S rRNA gene using the universal vertebrate primers Vert01 (Forward: 5’-ACACCGCCCGTCACTCT; Reverse: 5’-CTTCCGGTACACTTACCATG; De Barba et al. 2014;

Taberlet et al. 2018). For the Svalbard dataset, amplification targeted a ∼260 bp fragment of the mitochondrial 16S rRNA gene using the universal vertebrate primers Vert16S (Forward: 5’-AGACGAGAAGACCCYDTGGAGCTT; Reverse: 5’-GATCCAACATCGAGGTCGTAA; Vences et al. 2016). Each selected MOTU represented a particular taxonomic assignment case. MOTUs from the Guadeloupe dataset were named “Guad_port_complet_x” and MOTUs from the Svalbard dataset were named “MUSA_Vert16S_x”, where “x” corresponds to the MOTU identifier in the original dataset. For each dataset, several representative taxonomic assignment cases were presented step-by-step in order to illustrate how different situations are handled within the TRIDENT workflow.

The second test series was based on the same Guadeloupe and Svalbard datasets described above and consisted of re-analysing complete eDNA datasets that had previously been manually processed using similar curation and assignment criteria. This comparison aimed to evaluate the time efficiency of TRIDENT relative to the manual workflow while assessing the consistency of the resulting taxonomic assignments. For the Svalbard dataset, sequence-level taxonomic assignments produced by TRIDENT were compared with the original OBITools assignments (*obitag*; (Boyer et al. 2016), using the parameters described in Haderlé et al. (2026c), and with standard BLAST top-hit results obtained for the same MOTUs. The BLAST top-hit approach assigns each query sequence to the reference sequence in GenBank yielding the best alignment, based on the highest bit score. The comparison was used to determine whether the same taxa were recovered by the different approaches and, for discrepant cases, to identify the TRIDENT processing step responsible for the differences in taxonomic assignment.

The third test series was based on simulated datasets with known taxonomic assignments. These datasets were extracted from Bayer et al. (2025) and consisted of a realistic mock community derived from GBIF occurrence records of Actinopterygii and Chondrichthyes around Wadjemup (Rottnest Island, Western Australia). Simulated PCR amplicons targeting the 12S_Miya (Miya et al. 2015), 16S_Berry (Berry et al. 2017), and CO1_Leray (Leray et al. 2013) markers were extracted from available mitogenome sequence data. Duplicate or highly similar sequences were intentionally retained in order to better reflect realistic eDNA datasets (Bayer et al. 2025). Five amplicon sequencing libraries were subsequently simulated using ART v2.5.8 (Huang et al. 2012) to reproduce realistic Illumina sequencing error profiles for amplicon datasets. Using these simulated datasets, we compared the taxonomic assignment levels produced by TRIDENT with the known species-level assignments of the ASVs included in the simulated communities for the three barcode markers (101 ASVs for 12S, 108 ASVs for 16S, and 114 ASVs for CO1). Resulting graphs were produced using R version 4.4.3 (R Core Team 2025). In addition, for the 12S marker, the performance of each successive TRIDENT filtering stage was assessed by calculating precision, recall, and false rejection rate (FRR), based on the retention of the known reference species among the candidate species. Precision was calculated as the proportion of retained candidate species corresponding to the known reference species, while recall was calculated as the proportion of ASVs for which the correct reference species was retained. The FRR was calculated as the proportion of ASVs for which the correct reference species was no longer retained at a given filtering stage.

## RESULTS

### 1. Benchmarking Representative Assignment Cases

#### a. Two straightforward examples of high-confidence species-level assignments

MUSA_Vert16S_00005: BLAST searches against GenBank returned a single molecular match in the MOL list, *Odobenus rosmarus* (walrus), with a barcoding gap (**Fig. 2**) of 10.52 % to the closest related species (*Arctocephalus forsteri*). Querying WoRMS did not add additional taxa, as *Odobenus* is a monospecific genus. Geographic filtering using GBIF occurrence data, centered at latitude 78.5° N and longitude 14.5° E with a 250 km radius (Appendix 2), confirmed that *O. rosmarus* is well represented in the selected area (1,997 local GBIF occurrences on 11-02-2026). As no other plausible species was identified, no additional validation was required. TRIDENT therefore assigned this MOTU to *O. rosmarus* with high confidence.

**Figure 2:**
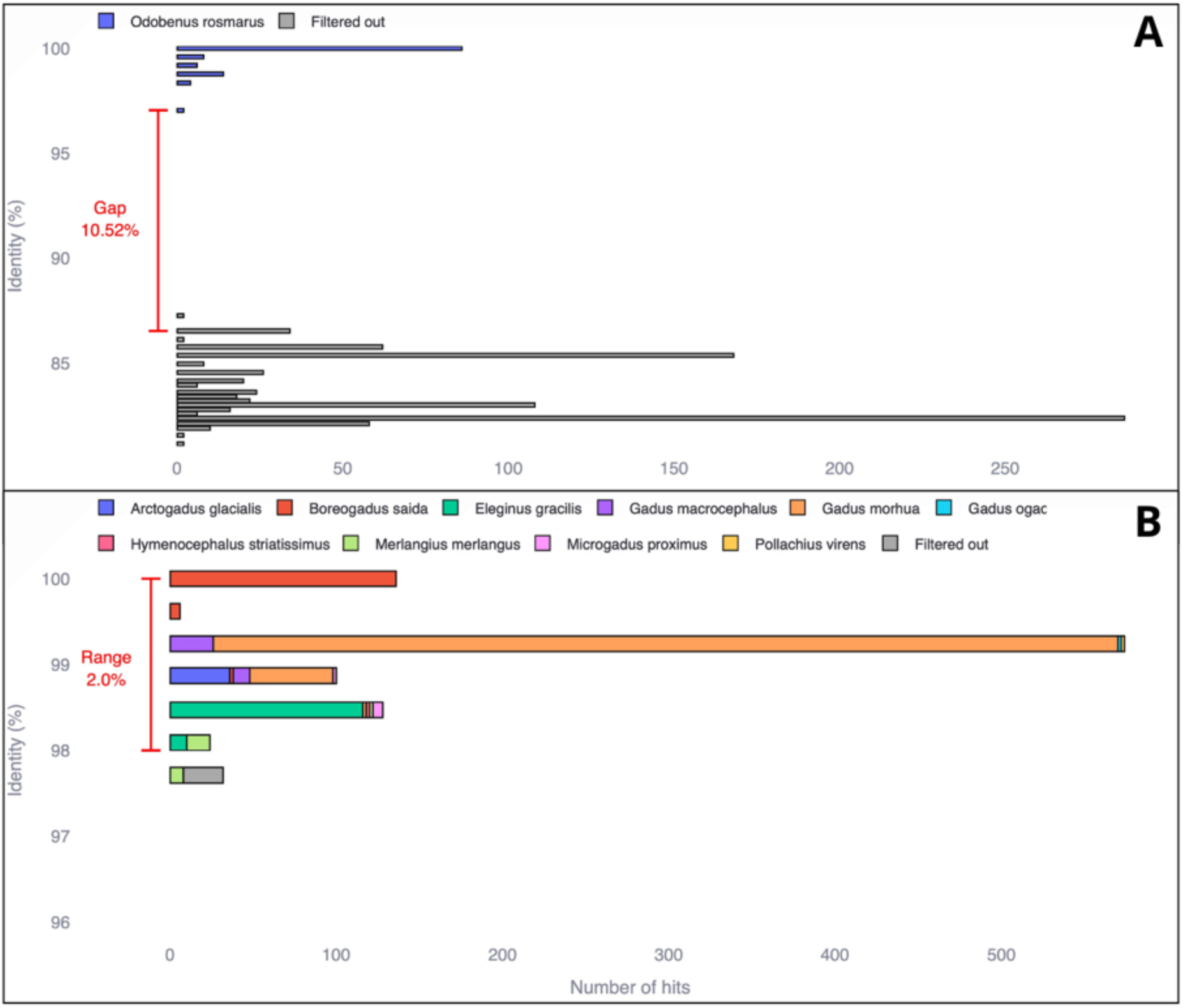
Top NCBI BLAST hits for two MOTUs in TRIDENT, plotted by percent identity and colored by matched species (grey for filtered out); MUSA_Vert16S_00005 (A) shows a clear barcoding gap (10.52% drop) isolating Odobenus rosmarus; MUSA_Vert16S_00013 (B) lacks such a gap, so assignment falls back to a similarity criterion.

Guad_port_complet_319: BLAST searches against GenBank returned five molecular matches in the MOL list, *Pterois volitans, P. miles, P. lunulata, Dendrochirus bellus and D. barberi*, with a barcoding gap of 3.03% to the closest related species (*Brachypterois serrulata*). Querying WoRMS added 11 additional species of *Pterois*. Geographic filtering using GBIF occurrence data, centered at latitude 16.20° N and longitude -61.5° E with a 150 km radius, informed that only *P. volitans* is well represented in the selected area (590 local GBIF occurrences on 03-24-2026). As no other plausible species was identified, no additional validation was required. TRIDENT therefore assigned this MOTU to *P. volitans* with high confidence.

#### b. Cases of taxonomic assignment to multiple plausible species

MUSA_Vert16S_00013: The initial GenBank blast search yielded 10 candidate species (with similarity values ranging from 100 to 98%; **Fig. 2**) across eight genera in the MOL list (*Arctogadus glacialis*, *Boreogadus saida*, *Eleginus gracilis*, *Gadus macrocephalus*, *G. morhua*, *G. ogac*, *Hymenocephalus striatissimus*, *Merlangius merlangus*, *Microgadus proximus*, *Pollachius virens*). The number of species was reduced to nine in the Sankey diagram (**Fig. 3**) following taxonomic harmonisation, which consolidated synonymous taxonomic names.Taxonomic inventory via WoRMS increased this list to 40 species. Applying the same geographic filter as above reduced the candidates to five locally distributed species: *A. glacialis* (34 GBIF occurrences), *B. saida* (2474occurrences), *G. morhua* (14,973 occurrences), *P. virens* (179 occurrences) and *M. merlangus* (12 occurrences). As all these five remaining species were already represented in GenBank, no further BOLD-based validation was performed. TRIDENT assigned this MOTU to five hypothetical species of the family Gadidae, all consistent with both molecular and geographic evidences (**Fig. 3**).

**Figure 3:**
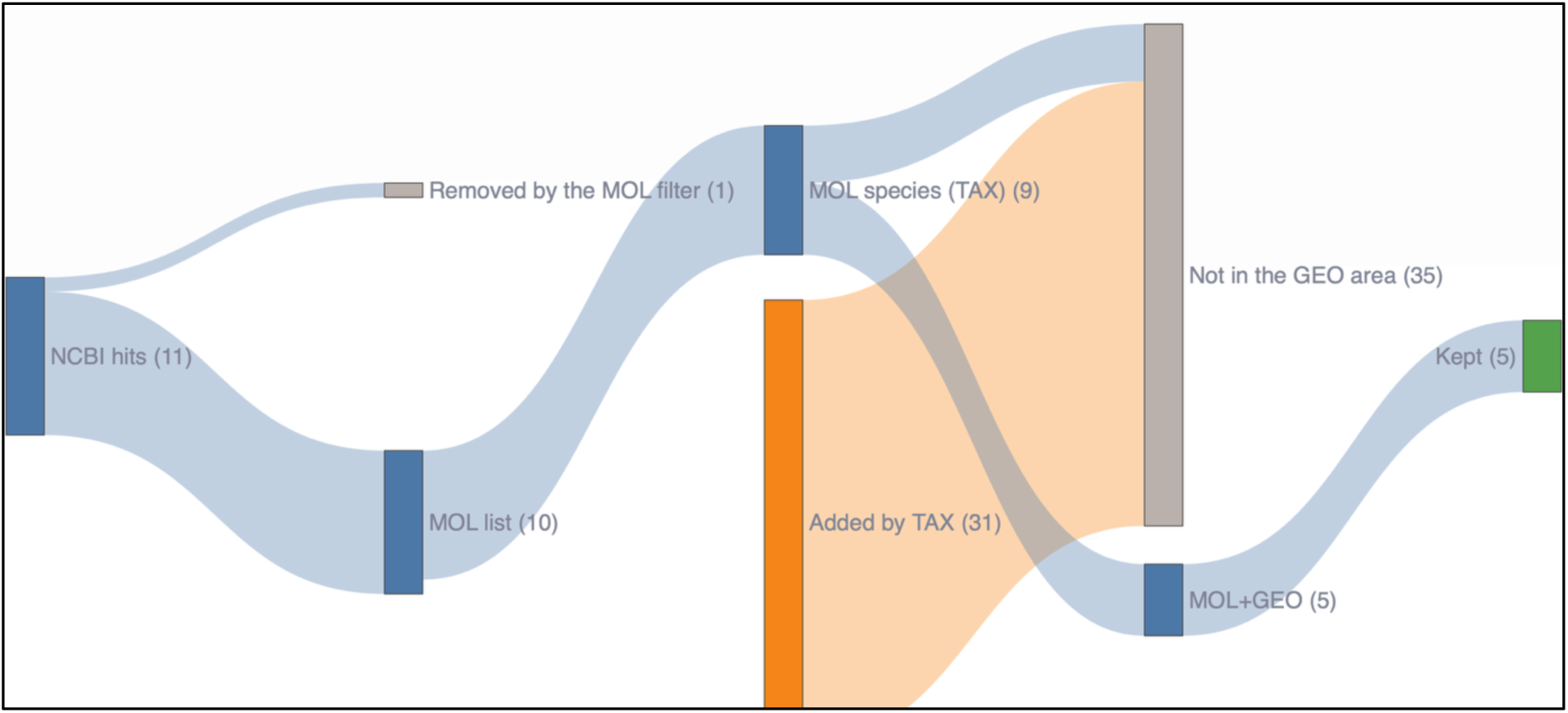
Sankey diagram illustrating the identification workflow for MOTU MUSA_Vert16S_00013. The diagram shows the number of candidate species retained or excluded at each successive filtering step. The initial molecular (MOL) step retained 10 species, nine of which were also retained after taxonomic (TAX) filtering, which additionally introduced 31 potential species. Geographic (GEO) filtering reduced the candidate list to five species, all originating from the initial MOL list. Following the final validation step, five hypothetical species were retained for taxonomic assignment.

Guad_port_complet_30: The initial GenBank blast search yielded five candidate species in the MOL list (*Ablennes hians, Tylosurus acus, T. crocodilus, T. melanotus and T. rafale*), with a 2% divergence from the closest species (*Strongylura incisa*). Taxonomic inventory of the two genera via WoRMS added six *Tylosurus* sp. Applying the same geographic filter as above reduced the candidates to two locally distributed species: *A. hians* (28 GBIF occurrences) and *T. crocodilus* (55 occurrences). As both species were represented in GenBank, no BOLD-based validation was performed. TRIDENT assigned this MOTU to two hypothetical species of the family Belonidae, both consistent with molecular and geographic evidences.

#### c. Identification of a probable taxon despite the absence of its barcode in GenBank

MUSA_Vert16S_00111: The initial GenBank search returned five species (similarity) in the MOL list (*T. forficatus*, *T. murrayi*, *T. pingelii*, *T. scepticus* and *Icelus spiniger,* with a maximum similarity of 98.47%). Taxonomic analysis via WoRMS added 20 additional *species*. Geographic filtering using GBIF occurrences retained two species of the MOL List present in the study area, *T. murrayi* (800 occurrences) and *T. pingelii* (135 occurrences), as well as *T. nybelini* and *I. bicornis*, added at the WoRMS step in the TAX list and supported respectively by 188 and 244 local occurrences. Because *T. nybelini* and *I. bicornis* were absent from the initial MOL list, TRIDENT looked for available CO1 barcode sequences for these species from BOLD and compared them to those of the closest molecular matches. Sequence similarities exceeded the similarity threshold (e.g. 98% identity with *T. pingelii* for *T. nybelini*), supporting molecular consistency among these taxa. A final check step confirmed the absence of reference barcode sequences for *T. nybelini* and *I. bicornis* in GenBank for the targeted marker. TRIDENT therefore assigned this MOTU to the Cottidae family, retaining four hypothetical species (*T. murrayi*, *T. pingelii*, *T. nybelini*, and *I. bicornis*; **Fig. 4**).

**Figure 4:**
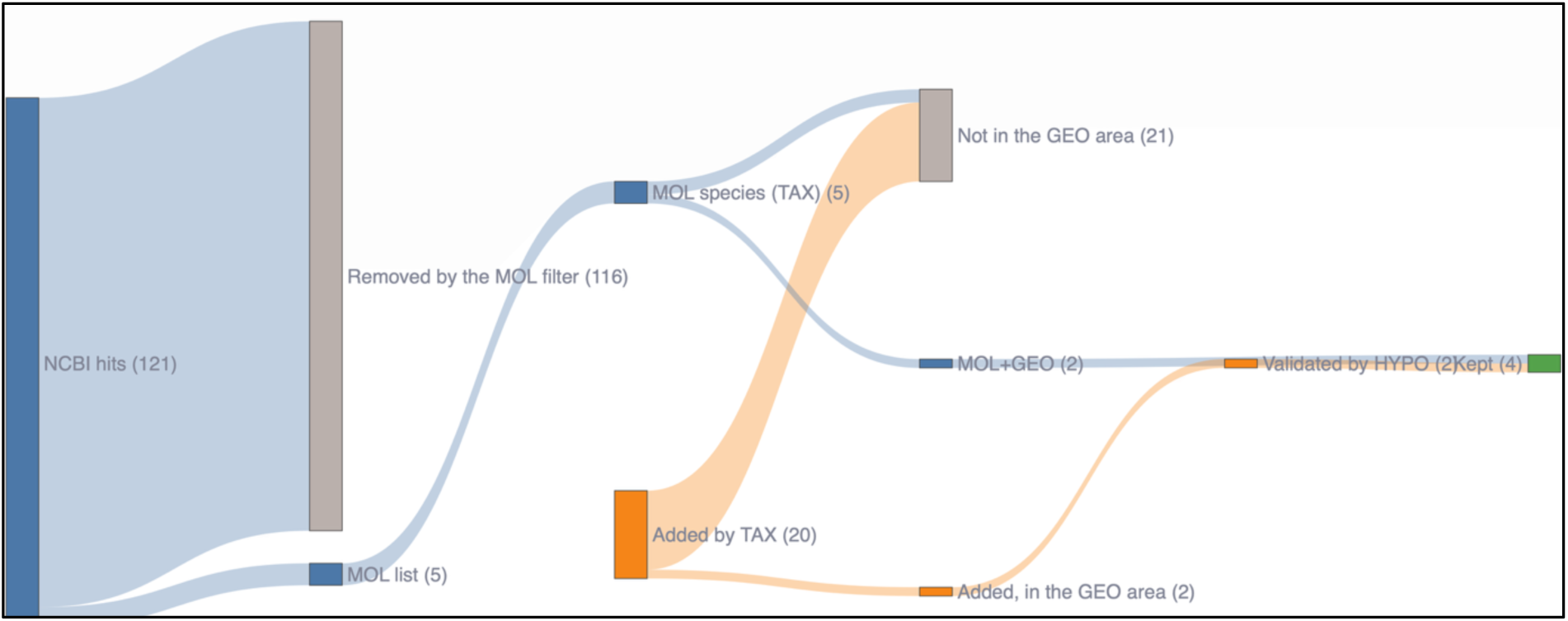
Sankey diagram illustrating the identification workflow for MOTU MUSA_Vert16S_00111. The initial molecular (MOL) step retained five candidate species, while the taxonomic (TAX) step added 20 additional species. Geographic (GEO) filtering retained two species from the initial MOL list and two species newly introduced at the TAX step. Both additional species were subsequently validated through comparison with COI barcode sequences available in BOLD and verification of their absence from GenBank. The final list comprised four hypothetical species retained for taxonomic assignment.

#### d. Detection of an artifact

MUSA_Vert16S_00065: GenBank searches returned three candidate taxa (*Balaena mysticetus, Balaenoptera borealis* and *Orcinus orca*), but with low sequence similarity (max. similarity 91.60%), below the default acceptance threshold. The following steps of TRIDENT showed that no additional species from the same genera were geographically plausible and absent from GenBank. Consequently, TRIDENT did not produce a taxonomic assignment for this MOTU.

Additional BLAST searches were conducted against the RefSeq Genome database (NCBI), restricting query to the organism *Delphinapterus leucas* (taxid:9749). These analyses revealed 100% similarity with a genomic contig of *D. leucas* (accession NW_022098063). As this species was independently detected in the same sample, the result indicates the amplification of a NUMT. This case highlights TRIDENT’s ability to conservatively reject unreliable assignments. in the presence of conflicting or insufficient molecular evidence.

### 2. Empirical Application to Complete eDNA Datasets

To assess the performance of TRIDENT relative to the fully manual taxonomic assignment workflow, we reanalyzed complete eDNA datasets that had previously been processed manually using the same successive criteria. For the Svalbard dataset (50 MOTUs in the FASTA file), TRIDENT completed the analysis in 30 min and identified a total of 33 distinct taxa, of which 22 were assigned to the species level with high confidence. In comparison, the same analysis performed manually required approximately 15 hours of full-time work and yielded an identical number of identified taxa.

For the GPMG Guadeloupe dataset (412 MOTUs in the initial FASTA file), TRIDENT completed the analysis in 3 h 25 min, resulting in 140 distinct teleost taxa. The equivalent manual workflow required approximately 40 hours of full-time effort to complete the taxonomic verification and assignment.

The comparison of BLAST top-hit and obitag taxonomic assignments with TRIDENT results for the Svalbard dataset (**Table 1**) revealed identical taxonomic assignments for 16 of the 50 MOTUs. For the remaining MOTUs, discrepancies could be attributed to several steps of the TRIDENT workflow. Twelve MOTUs were removed at the taxonomic (TAX) step, which retains only marine species (WoRMS): ten corresponded to potential contaminants (human and domestic or laboratory animals) and two to local terrestrial species (*Rangifer tarandus* and *Branta leucopsis*). Three additional MOTUs, all assigned to *Phocoena phocoena* by obitag, showed low best-hit identity (87.9–95.4%) and were identified as suspected nuclear mitochondrial DNA segments (NUMTs; 100% match to a beluga genome contig), so that TRIDENT returned no species-level assignment for them. Five MOTUs were resolved to a single species by the geographic (GEO) filter, which discarded the non-local species (mostly Pacific) tied with the top hit. Thirteen further MOTUs were assigned at the genus, family or order level, mainly because of the absence of a clear barcode gap, which limited taxonomic resolution at the species level, or because of incomplete reference database coverage (local species with no 16S sequence in GenBank, added by the HYPO step). In such cases, TRIDENT favoured more conservative taxonomic assignments rather than potentially unreliable species-level identifications. Finally, one apparent discrepancy (*Phoca groenlandica* vs *Pagophilus groenlandicus*) resulted from differences in taxonomic nomenclature rather than from an actual disagreement in species identification.

**Table 1:** Comparison of taxonomic assignments obtained using obitag, BLAST top-hit identification, and the TRIDENT workflow for the Svalbard dataset. The table presents the MOTU identifiers (Seq_ID), taxonomic assignments generated by each approach, and explanations for discrepancies between methods

| Seq_ID | obitag | BLAST top hit | TRIDENT | Reason for the difference |
|---|---|---|---|---|
| MUSA_Vert16S_00001 | *Homo sapiens* | *Homo sapiens* (100%) | No species | Not a marine species: removed at the TAX step (WoRMS, marine species only) |
| MUSA_Vert16S_00004 | *Delphinapterus leucas* | *Delphinapterus leucas* (100%) | *Delphinapterus leucas* | Identical assignment |
| MUSA_Vert16S_00005 | *Odobenus rosmarus* | *Odobenus rosmarus* (100%) | *Odobenus rosmarus* | Identical assignment |
| MUSA_Vert16S_00006 | *Myoxocephalus* | *Myoxocephalus aenaeus, Myoxocephalus ochotensis, Myoxocephalus scorpius* (100%) | *Myoxocephalus scorpius* | Tied 100% hits with Pacific sculpins; only *M. scorpius* is local (GEO) |
| MUSA_Vert16S_00008 | *Clupea harengus* | *Clupea harengus* (100%) | *Clupea harengus* | Identical assignment |
| MUSA_Vert16S_00011 | *Cyclopterus lumpus* | *Cyclopterus lumpus* (100%) | *Cyclopterus lumpus* | Identical assignment |
| MUSA_Vert16S_00012 | Pleuronectidae | *Cleisthenes herzensteini, Cleisthenes pinetorum, Hippoglossoides dubius* (+4) (100%) | *Hippoglossoides platessoides* | Tied 100% hits with mostly Pacific flatfish; only *H. platessoides* is local (GEO) |
| MUSA_Vert16S_00013 | *Boreogadus saida* | *Boreogadus saida* (100%) | Gadidae (family), 5 species | Several local species within the similarity range (no barcoding gap): 5 local cods |
| MUSA_Vert16S_00014 | *Uria lomvia* | *Uria lomvia* (98.8%) | *Uria lomvia* | Identical assignment |
| MUSA_Vert16S_00016 | Somateria | *Somateria mollissima, Somateria spectabilis* (100%) | Anatidae (family), 4 species | Several local species within the similarity range (no barcoding gap): 3 Somateria and *Polysticta stelleri* |
| MUSA_Vert16S_00021 | *Gymnocanthus tricuspis* | *Gymnocanthus tricuspis* (100%) | *Gymnocanthus tricuspis* | Identical assignment |
| MUSA_Vert16S_00022 | *Rangifer tarandus* | *Rangifer tarandus* (100%) | No species | Not a marine species: removed at the TAX step (WoRMS, marine species only); a local land mammal |
| MUSA_Vert16S_00023 | Lumpeninae | *Leptoclinus maculatus, Lumpenus fabricii, Lumpenus lampretaeformis* (100%) | Lumpenidae (family), 4 species | Several local species within the similarity range (no barcoding gap): 4 local pricklebacks |
| MUSA_Vert16S_00024 | *Gallus* | *Gallus gallus, Gallus sonneratii* (100%) | No species | Not a marine species: removed at the TAX step (WoRMS, marine species only) |
| MUSA_Vert16S_00025 | *Liparis fabricii* | *Liparis fabricii* (99.62%) | *Liparis* sp., 4 species | Several local species within the similarity range (no barcoding gap); HYPO adds *L. tunicatus* (no 16S in GenBank) |
| MUSA_Vert16S_00028 | *Anarhichas* | *Anarhichas lupus, Anarhichas minor* (100%) | *Anarhichas* sp., 3 species | Several local species within the similarity range (no barcoding gap): 3 local *Anarhichas* |
| MUSA_Vert16S_00030 | *Mallotus villosus* | *Mallotus villosus* (99.25%) | *Mallotus* sp., 2 species | HYPO adds *M. catervarius* (no 16S in GenBank, COI identical to *M. villosus*) |
| MUSA_Vert16S_00031 | *Bos taurus* | *Atherurus africanus, Bos grunniens, Bos indicus* (+2) (100%) | No species | Not a marine species; the marine relatives of a mislabelled hit (*Phascolosoma*) have no local GBIF record (GEO) |
| MUSA_Vert16S_00032 | *Phoca vitulina* | *Phoca vitulina* (100%) | Phocidae (family), 2 species | Several local species within the similarity range (no barcoding gap): *Phoca vitulina* and *Pusa hispida* (98.4%) |
| MUSA_Vert16S_00033 | *Canis lupus* | *Canis lupus* (100%) | No species | Not a marine species: removed at the TAX step (WoRMS, marine species only) |
| MUSA_Vert16S_00035 | *Alle alle* | *Alle alle* (100%) | *Alle alle* | Identical assignment |
| MUSA_Vert16S_00040 | *Melanogrammus aeglefinus* | *Melanogrammus aeglefinus* (100%) | *Melanogrammus aeglefinus* | Identical assignment |
| MUSA_Vert16S_00041 | *Sus* | *Sus scrofa* (100%) | No species | Not a marine species: removed at the TAX step (WoRMS, marine species only) |
| MUSA_Vert16S_00042 | *Branta leucopsis* | *Branta leucopsis* (100%) | No species | Not a marine species: removed at the TAX step (WoRMS, marine species only); its marine congeners have no local GBIF record (GEO). A local breeding bird |
| MUSA_Vert16S_00043 | *Cepphus grylle* | *Cepphus grylle* (100%) | *Cepphus grylle* | Identical assignment |
| MUSA_Vert16S_00044 | *Felis* | *Felis catus, Felis silvestris* (100%) | No species | Not a marine species: removed at the TAX step (WoRMS, marine species only) |
| MUSA_Vert16S_00045 | *Gasterosteus* | *Gasterosteus aculeatus, Gasterosteus gymnurus* (100%) | *Gasterosteus aculeatus* | Tied top hits with *G. gymnurus*; only *G. aculeatus* is local (GEO) |
| MUSA_Vert16S_00047 | *Ammodytes* | *Ammodytes marinus, Ammodytes* sp. (100%) | *Ammodytes marinus* | Top hit shared with an unnamed *Ammodytes* sp.; only *A. marinus* is local (GEO) |
| MUSA_Vert16S_00053 | *Anser brachyrhynchus* | *Anser brachyrhynchus* (100%) | *Anser* sp., 4 species | Several local species within the similarity range (no barcoding gap): 4 local *Anser* |
| MUSA_Vert16S_00065 | *Phocoena phocoena* | *Eubalaena australis* (91.6%) | No species | Best identity 91.6%, flagged as low identity; local cetaceans kept, no shared taxon below the order. A NUMT (beluga genome contig at 100%) |
| MUSA_Vert16S_00066 | *Vulpes lagopus* | *Vulpes lagopus (*100%) | *Vulpes lagopus* | Identical assignment |
| MUSA_Vert16S_00070 | *Phocoena phocoena* | *Balaenoptera borealis* (87.87%) | No species | Best identity 87.9%, flagged as low identity; local cetaceans kept, no shared taxon below the order. A NUMT (beluga genome contig at 100%) |
| MUSA_Vert16S_00073 | *Erignathus barbatus* | *Erignathus barbatus* (99.59%) | *Erignathus barbatus* | Identical assignment |
| MUSA_Vert16S_00074 | *Phoca groenlandica* | *Phoca groenlandica* (99.59%) | *Pagophilus groenlandicus* | Same species; current WoRMS name |
| MUSA_Vert16S_00077 | *Phocoena phocoena* | *Balaenoptera borealis* (95.4%) | No species | Best identity 95.4%, flagged as low identity; local cetaceans kept, no shared taxon below the order. A NUMT (beluga genome contig at 100%) |
| MUSA_Vert16S_00078 | *Ovis* | *Ovis aries, Ovis orientalis* (100%) | No species | Not a marine species: removed at the TAX step (WoRMS, marine species only) |
| MUSA_Vert16S_00082 | *Homo sapiens* | *Homo sapiens, Mus musculus* (99.2%) | No species | Not a marine species: removed at the TAX step (WoRMS, marine species only) (hits on human and mouse nuclear clones) |
| MUSA_Vert16S_00085 | *Amblyraja radiata* | *Amblyraja radiata* (100%) | *Amblyraja* sp., 2 species | HYPO check finds the 16S of *A. hyperborea* at 98.9%, which the MOL hit limit had hidden |
| MUSA_Vert16S_00093 | *Sterna paradisaea* | *Sterna paradisaea* (100%) | *Sterna paradisaea* | Identical assignment |
| MUSA_Vert16S_00095 | *Salvelinus* | *Salvelinus alpinus, Salvelinus drjagini, Salvelinus umbla* (100%) | *Salvelinus alpinus* | Tied 100% hits with 32 chars and whitefish; only *S. alpinus* is local (GEO) |
| MUSA_Vert16S_00110 | *Rissa tridactyla* | *Rissa tridactyla* (100%) | *Rissa tridactyla* | Identical assignment |
| MUSA_Vert16S_00111 | Cottidae | *Icelus spiniger, Triglops forficatus, Triglops pingelii* (+1) (98.47%) | Cottidae (family), 4 species | Top hit on a Pacific *Icelus*; 2 local sculpins in the similarity range, and HYPO adds *T. nybelini* and *I. bicornis* (no 16S in GenBank) |
| MUSA_Vert16S_00117 | *Cavia porcellus* | *Cavia porcellus* (100%) | No species | Not a marine species: removed at the TAX step (WoRMS, marine species only) |
| MUSA_Vert16S_00130 | *Icelus* | *Icelus spatula* (98.47%) | Cottidae (family), 5 species | Local *Icelus* and *Triglops* in the similarity range; HYPO adds *T. nybelini* and *I. bicornis* |
| MUSA_Vert16S_00138 | *Eumicrotremus spinosus* | *Eumicrotremus spinosus* (100%) | *Eumicrotremus spinosus* | Identical assignment |
| MUSA_Vert16S_00149 | *Cottales* | *Icelus spiniger* (96.56%) | Cottidae (family), 5 species | Top hit on a Pacific *Icelus* (96.6%); local sculpins in the similarity range; HYPO adds *T. nybelini* and *I. bicornis* |
| MUSA_Vert16S_00162 | *Meleagris gallopavo* | *Labidesthes sicculus, Meleagris gallopavo* (100%) | No species | Not a marine species: removed at the TAX step (WoRMS, marine species only) (the freshwater *Labidesthes* hit is a mislabelled record) |
| MUSA_Vert16S_00181 | *Cottales* | *Icelus spiniger* (95.8%) | Perciformes (order), 6 species | Top hit on a Pacific *Icelus* (95.8%); local sculpins and the lumpsucker in the similarity range; HYPO adds *T. nybelini* and *I. bicornis* |
| MUSA_Vert16S_00188 | *Leptoclinus maculatus* | *Leptoclinus maculatus* (99.62%) | *Leptoclinus maculatus* | Identical assignment |
| **MUSA_Vert16S_00195** | *Balaenoptera acutorostrata* | *Balaenoptera acutorostrata (*99.58%) | *Balaenoptera acutorostrata* | Identical assignment |

## TRIDENT (Taxonomic Resolution and IDentification using Environmental dNa Traces)

### 3. Application to Simulated eDNA Datasets

For the 12S marker, 66% of species were assigned by TRIDENT to the species level (**Fig. 5**). In 27% of cases, assignments were downgraded to the genus level, mainly due to incomplete reference databases (more than half of the cases) or because multiple species shared the same assignment profile (representing over 40% of the cases). Assignments were downgraded to the family level in 4% of cases and to the order level in 1% of cases. Finally, 2% of ASVs remained unassigned, because no corresponding occurrence records were available in GBIF for the expected species.

**Figure 5:**
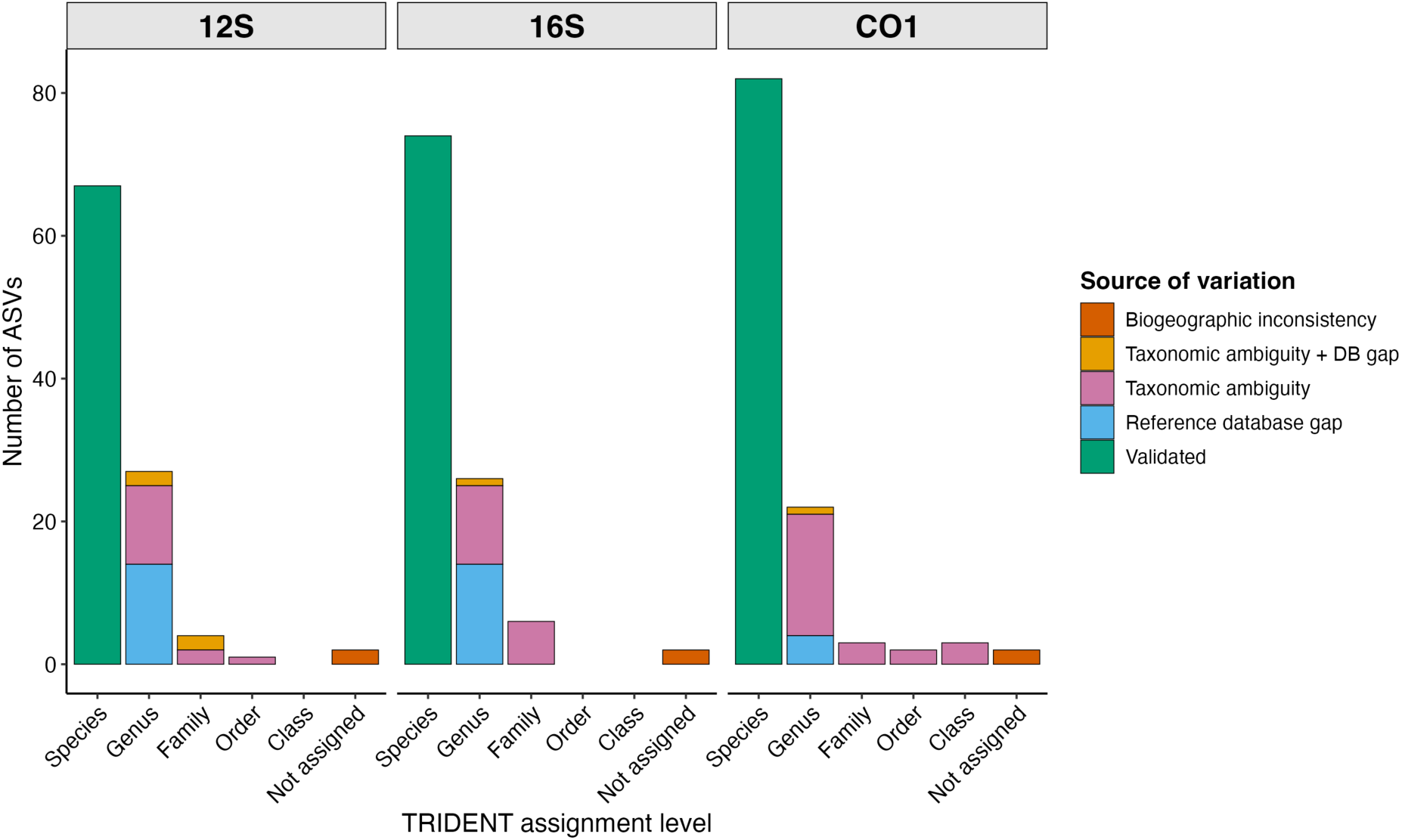
Stacked bar plots showing the taxonomic assignment level reached by TRIDENT for the simulated dataset of Bayer et al. (2025), in which all ASVs are assigned at species level. Each panel represents a primer set, bars show the number of ASVs per TRIDENT assignment level (species, genus, family, order, class, or not assigned), and colors denote assignment outcomes: Validated (green), identical assignment; Reference database gap (blue), presence of at least one closely related species lacking barcode data in GenBank; Taxonomic ambiguity (pink), inability to discriminate among molecularly similar taxa; Taxonomic ambiguity + database gap (orange), combined effect of ambiguity and incomplete reference data; and Biogeographic inconsistency (red), assignment rejected because GBIF occurrence thresholds were not met within the target geographic region.

For the 16S marker, 68.5% of species were assigned to the correct species level (**Fig. 5**), a result very similar to that obtained with 12S. Assignments were downgraded to the genus level in 24% of cases, mainly due to incomplete reference databases (54% of cases) or the presence of multiple candidate species (more than 42% of cases). Family-level assignments represented 6% of cases, while less than 2% of ASVs remained unassigned.

For the CO1 marker, 72% of species were assigned at the species level (**Fig. 5**). Assignments were downgraded to the genus level in slightly more than 19% of cases, and approximately 2% of ASVs remained unassigned, values comparable to those observed for the 12S and 16S markers. However, CO1 was the only marker for which some ASVs were downgraded to a taxonomic rank as high as the class level (representing around 3% of ASVs).

The performance of the TRIDENT identification pipeline was evaluated at each filtering stage using precision, recall and false rejection rate (FRR), based on 12S ASVs **(Appendix 3**). At the MOL stage, precision was 35%, while recall reached 100%, indicating that all reference species were retained among the candidate species, although a relatively large number of additional candidates remained. The addition of taxonomic filtering (MOL + TAX) resulted in a slight improvement in precision (37%), without affecting recall (100%). The inclusion of geographic filtering (MOL + TAX + GEO) substantially improved precision, which increased to 67.16%. However, four reference species were no longer retained among the candidates because their occurrence records in GBIF did not meet the geographic filtering thresholds. This resulted in a decrease in recall to 95.74% and a false rejection rate of 4.26%. At the FINAL stage, precision decreased to 61.22%, while recall and FRR remained unchanged. This decrease in precision was due to the reintroduction of additional candidate species lacking barcode sequences in GenBank, which were therefore added to the list of hypothetical candidate species.

## DISCUSSION

We present TRIDENT, an optimized tool that improves taxonomic assignments of marine vertebrates in eDNA metabarcoding by integrating molecular, taxonomic, and biogeographical evidence. Its performance is assessed through case studies and complete eDNA datasets.

### 1. Reaffirming the Role of Taxonomy in eDNA-based Biomonitoring

Over the past decade, DNA-based bioassessment has explored taxonomy-free approaches, OTU-based indices, machine-learning pipelines (Cordier et al. 2018; Tapolczai et al. 2019), and phylogenetic inference from sequence clusters (Keck et al. 2018). These methods work well for microorganisms and micro-eukaryotes with poorly resolved species boundaries (Feio et al. 2020; Gregersen et al. 2023), but their application to macro-organisms, particularly vertebrates, remains unsatisfactory, as conservation policy, legislation, and management actions still rely on species-level information (Maclaurin and Sterelny 2008).

In this context, TRIDENT’s primary contribution is to place molecular taxonomy back at the center of eDNA metabarcoding studies. Molecular taxonomy requires both expertise in molecular biology and a solid understanding of the principles of taxonomy. Many studies on environmental DNA are sometimes conducted by scientists who have not received in-depth training in either of these two fields, or even both. This increases the risk that the molecular taxonomy stage of the process will not be optimal, and that automated assignments are accepted uncritically. TRIDENT is designed as an assistant to the molecular taxonomist, highly experienced or less, providing a structured and transparent framework to evaluate, refine, and justify taxonomic assignments.

A key design principle of TRIDENT is user-controlled flexibility. Different parameters can be adjusted, at each step, to accommodate different taxonomic contexts, ecological backgrounds, geographic scales, and data availability scenarios.

### 2. Performance Evaluation of TRIDENT

The benchmarking of specific case studies presented in this work highlights the behaviour of TRIDENT in addressing common scenarios that complicate taxonomic assignment in eDNA metabarcoding. Overall, the results demonstrate the robustness of the TRIDENT framework across both empirical and simulated datasets. Across all comparisons, taxonomic assignments were fully concordant at higher taxonomic ranks, meaning that when species-level identification was not possible, the resulting assignments remained consistent with the expected taxonomy.

Consistent performance was observed across the different molecular markers tested. This highlights an important feature of TRIDENT: its gene-agnostic design, which allows its application to vertebrate metabarcoding datasets regardless of the genetic marker or barcode used. Unlike approaches specifically developed around a single barcode, TRIDENT relies on a modular framework that combines molecular, taxonomic, and biogeographical information, making it adaptable to different marker-specific reference datasets and metabarcoding applications. Simulated datasets were deliberately included in this study because, despite repeated recommendations for their use in methodological validation (Gardner et al. 2019), they remain underutilized in metabarcoding research. In contrast to real environmental samples, where the true species composition is unknown, simulated communities provide a controlled framework that enables direct evaluation of taxonomic assignment accuracy and the identification of methodological limitations (O’Rourke et al. 2020; Lotterhos et al. 2022). Such benchmarking approaches are well established in fields such as machine learning (Thiyagalingam et al. 2022) and are increasingly recognized as essential for the rigorous evaluation of eDNA classification methods (Bayer et al. 2025).

In their comparative study, Bayer et al. (2025) evaluated multiple datasets and nine taxonomic classifiers across three markers, reporting that, on average, classifiers correctly assigned species-level labels to 48% of ASVs. In comparison, TRIDENT achieved an average of 69% correct species-level assignments across the three markers tested, albeit on a subset of the dataset (the realistic simulated community). Although these results are not directly fully comparable due to differences in scope and dataset selection, they suggest an overall improvement in assignment performance when incorporating explicit molecular, taxonomic, and biogeographical constraints.

### 3. Alignment with FAIR Data Principles and Open-access Databases

The FAIR principles (Findable, Accessible, Interoperable, and Reusable) aim to improve the management and sharing of scholarly digital research objects, such as data and scripts (Wilkinson et al. 2016). Open sharing of eDNA data is essential for both research programs investigating global change impacts on biodiversity and programs supporting public policy, as it facilitates future reanalyses, long-term monitoring and the revision of assignments as reference databases and analytical tools improve. Although sharing complex eDNA datasets remains challenging, standardization efforts are developing, building on existing standards (Yilmaz et al. 2011; Wieczorek et al. 2012) and recent comprehensive guidelines on required metadata fields for eDNA studies (Takahashi et al. 2025).

To align with this movement toward standardization and open science, TRIDENT includes functionality to export standardized output tables that comply with FAIR data principles adapted to eDNA datasets (Wilkinson et al. 2016; Abarenkov et al. 2023; Takahashi et al. 2025). These tables include mandatory and recommended fields describing the taxonomic information associated with each (M)OTU/ASV occurrence, facilitating downstream data sharing and interoperability.

TRIDENT relies entirely on open-access databases (GenBank, GBIF, BOLD, WoRMS). While these resources may contain errors or taxonomic inconsistencies (Meiklejohn et al. 2019; Ribeiro et al. 2022), TRIDENT mitigates their impact through conservative assignment rules: it considers barcoding gaps, evaluates similarity across multiple closely related taxa when necessary, and requires a minimum number of occurrence records for geographic validation.

### 4. Preventing Misclassification and Data Loss

TRIDENT explicitly accounts for taxa absent from reference databases such as GenBank or BOLD. These databases are essential for eDNA taxonomic assignment, but coverage remains uneven across taxa, markers, and regions (Marques et al. 2021; Schmidt et al. 2025; Jung et al. In press), limiting species-level identification and slowing adoption in conservation (Beng and Corlett 2020). Expanding reference databases is essential but time-consuming and unevenly achievable. TRIDENT provides a complementary solution by identifying candidate taxa lacking molecular references and evaluating plausibility based on taxonomic similarity and regional species inventories. In regions with substantial molecular reference database gaps, this prevents potentially valid detections from being discarded or misclassified.

Geographic plausibility is incorporated using occurrence data from GBIF, helping to exclude ecologically implausible assignments, such as geographically distant taxa sharing similar barcodes. Although GBIF has known biases, the large data volume (>3.5 billion records) enables effective filtering when combined with user-defined thresholds for minimum occurrences. This flexibility enables adaptation to different ecological contexts, such as excluding pelagic taxa from coastal systems or broadening spatial filters in data-poor regions.

Contamination is also implicitly addressed. TRIDENT, developed and tested on vertebrate datasets, accounts for common contaminants such as *Homo sapiens*, domestic animals, or livestock species frequently associated with human activity or food sources.

Terrestrial taxa are removed at the WoRMS step, and any that remain are often excluded during the GBIF filtering step due to the lack of relevant occurrence records in natural ecosystems.

### 5. Limitations and Future Perspectives

The use of TRIDENT inevitably increases processing time relative to fully automated pipelines. However, this cost remains marginal compared to manual verification and substantially reduces the risk of misinterpretation, particularly with respect to species distributions. Taxa absent from GBIF-covered regions cannot be validated geographically, although TRIDENT explicitly flags such cases, allowing users to identify which validation step failed and to conduct targeted follow-up analyses. Furthermore, because TRIDENT allows users to adjust several parameters at each step of the process, it reflects the framework’s emphasis on molecular taxonomic expert-driven decision-making.

Regarding scalability, the main constraint lies in NCBI’s remote BLAST service rather than in TRIDENT itself. While the first three steps (MOL, TAX, GEO) can process datasets comprising up to several thousand (M)OTUs/ASVs, the HYPO step may require several days for larger datasets. To optimize processing time, we recommend splitting large datasets into separate FASTA files of approximately 100 sequences each and running them independently, particularly for the HYPO step. Processing can also be interrupted and resumed without losing completed searches.

Although developed and tested on marine vertebrate eDNA metabarcoding datasets, TRIDENT is not restricted to a specific genetic marker or barcode, and can therefore accommodate different marker-based metabarcoding approaches targeting vertebrates. Its modular architecture could also facilitate its adaptation to other taxonomic groups through the integration of additional reference databases, thereby extending its applicability beyond marine vertebrates. Future developments may include broader community testing, deployment as a server-based or cloud-accessible application to facilitate user access and simplify its use through a centralized interface, andintegration with downstream biodiversity analyses, such as the direct computation of diversity metrics from TRIDENT-validated taxonomic lists.

Overall, TRIDENT provides a robust, transparent, and expert-oriented framework for taxonomic validation in eDNA metabarcoding. By reconciling molecular evidence with taxonomic and ecological knowledge, it strengthens confidence in species-level inferences and enhances the reliability of eDNA-based biodiversity assessments.

## FUNDING ACKNOWLEDGMENT

This research received no specific funding.

## ACKNOWLEDGMENT

The authors would like to thank Lila Makouche, M.Sc. student in Bioinformatics, for her contribution to the development of the initial digital version of the protocol. We also thanks Erwan Quéméré and Émilien Foucaud (UMR DECOD, INRAE, Rennes France) who tested a first version of TRIDENT.

## DATA AVAILABILITY STATEMENT

The TRIDENT source code, documentation, and associated resources are available on GitHub at https://github.com/ISYEB-ADNe/TRIDENT. All datasets generated or analyzed during this study have been deposited in Dryad and are available to editors and reviewers via the following private reviewer link: http://datadryad.org/share/LINK_NOT_FOR_PUBLICATION/Ucno7YP83Ts1CNncoohFN68_LlZ30mgYjEUhHiKhziM. All scripts used for the analyses have been deposited in Zenodo (DOI: https://doi.org/10.5281/zenodo.21108395) and are accessible through the Dryad reviewer link.

## Appendix 1: Overview of the most commonly used nomenclatures for naming molecular units in eDNA metabarcoding

| Acronym | Full name | Definition | Purpose / context | Reference(s) |
|---|---|---|---|---|
| OTU | Operational Taxonomic Unit | Cluster of sequences grouped using a fixed similarity threshold (commonly 97%) | Early standardisation of HTS metabarcoding outputs, mainly in microbial ecology | Sneath & Sokal, 1973; Schloss & Handelsman, 2005 |
| MOTU | Molecular Operational Taxonomic Unit | OTU explicitly defined from molecular data, often applied across metazoans | Extension of OTU concept beyond microbiology | Blaxter et al., 2005 |
| ASV | Amplicon Sequence Variant | Exact biological sequences inferred after error correction; no clustering | High-resolution, reproducible units comparable across studies | Callahan et al., 2017 |
| ESV | Exact Sequence Variant | Functional synonym of ASV | Same as ASV | Callahan et al., 2016 |
| ZOTU | Zero-radius Operational Taxonomic Unit | OTU with zero clustering radius (100% identity), equivalent to ASVs (UNOISE) | Alternative denoising approach | Edgar, 2016 |

## Appendix 2: Screenshot of the third step of TRIDENT, dedicated to checking geographic consistency using GBIF occurrence data, showing an example of a geographic area delineated for searching taxon occurrences.

| Latitude | Longitude | Search extents (km) |
|---|---|---|
| 78.5 | 14.5 | 250 |
| Parsed as: 78.5000° | Parsed as: 14.5000° | Parsed extents: **250 km** |

## Appendix 3: Table used to calculate performance metrics (precision, recall and false rejection rate, FRR) at each filtering stage, 893 based on the 12S ASVs from Bayer et al. (2025). Each row corresponds to one ASV, and the TRUTH column gives the taxon to which that ASV belongs in the simulated dataset. The following columns give the number of taxa retained at each filtering stage. Columns with the suffix _correct are coded 1 when the TRUTH taxon was retained at the corresponding stage and 0 when it was not.

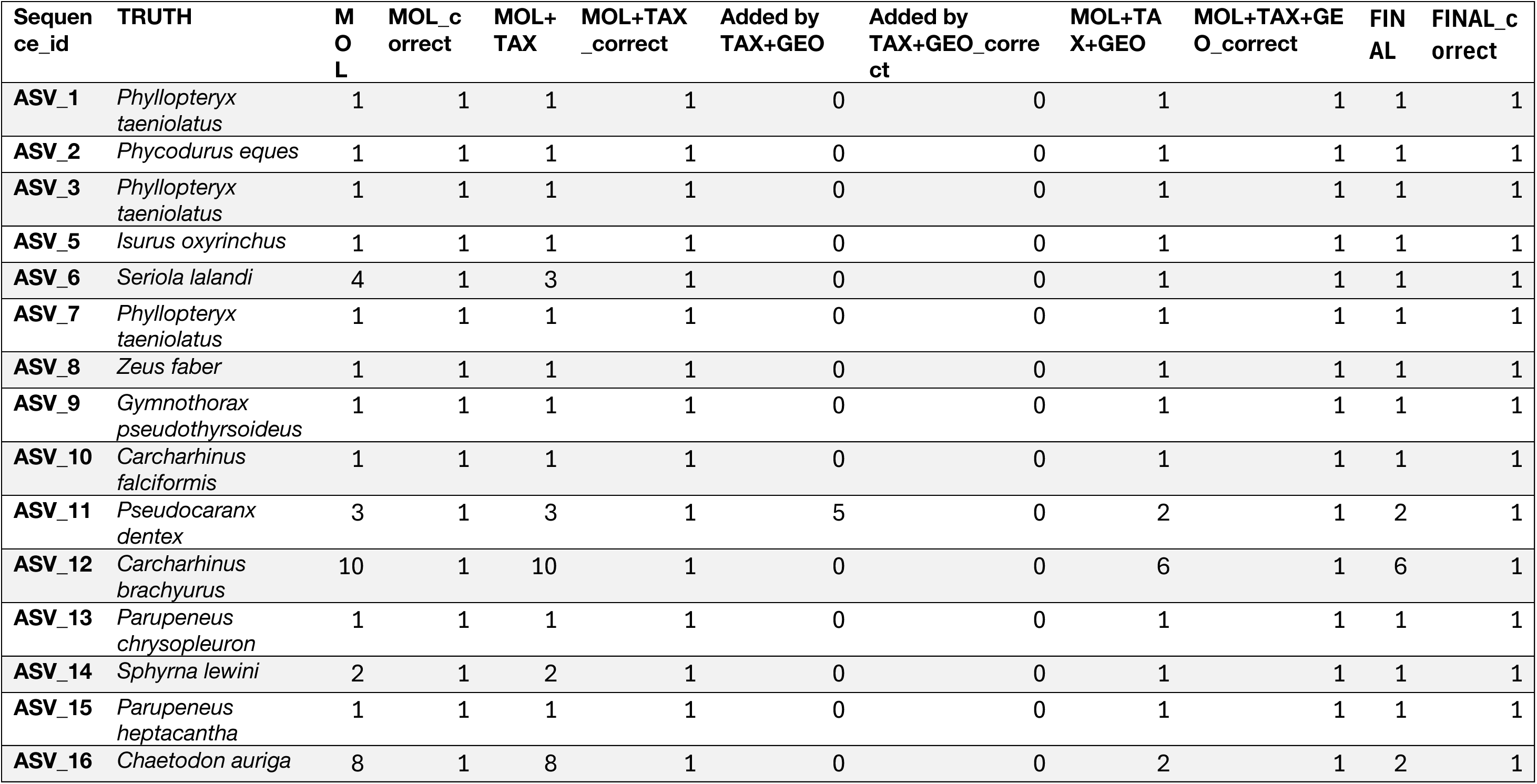
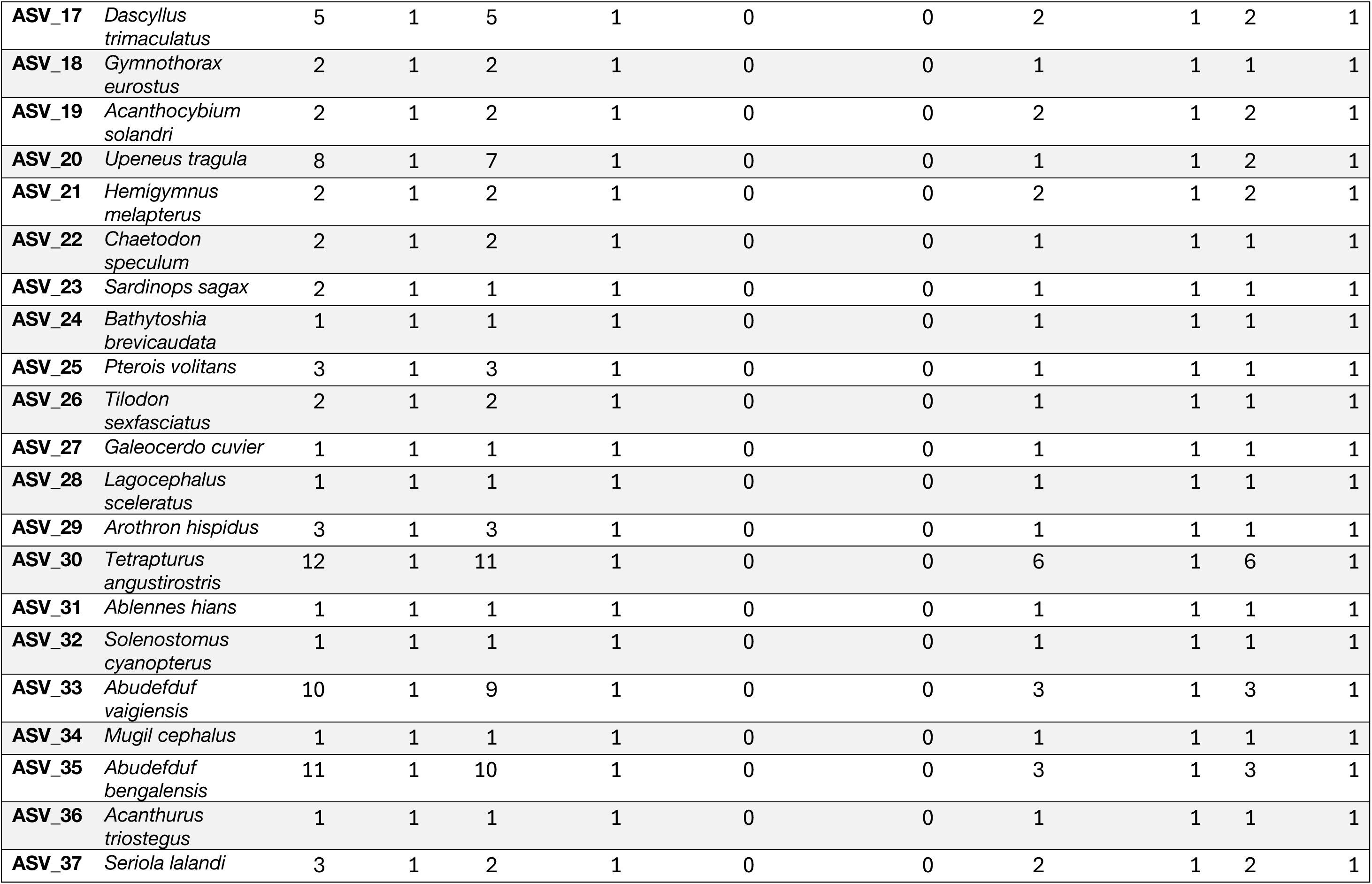
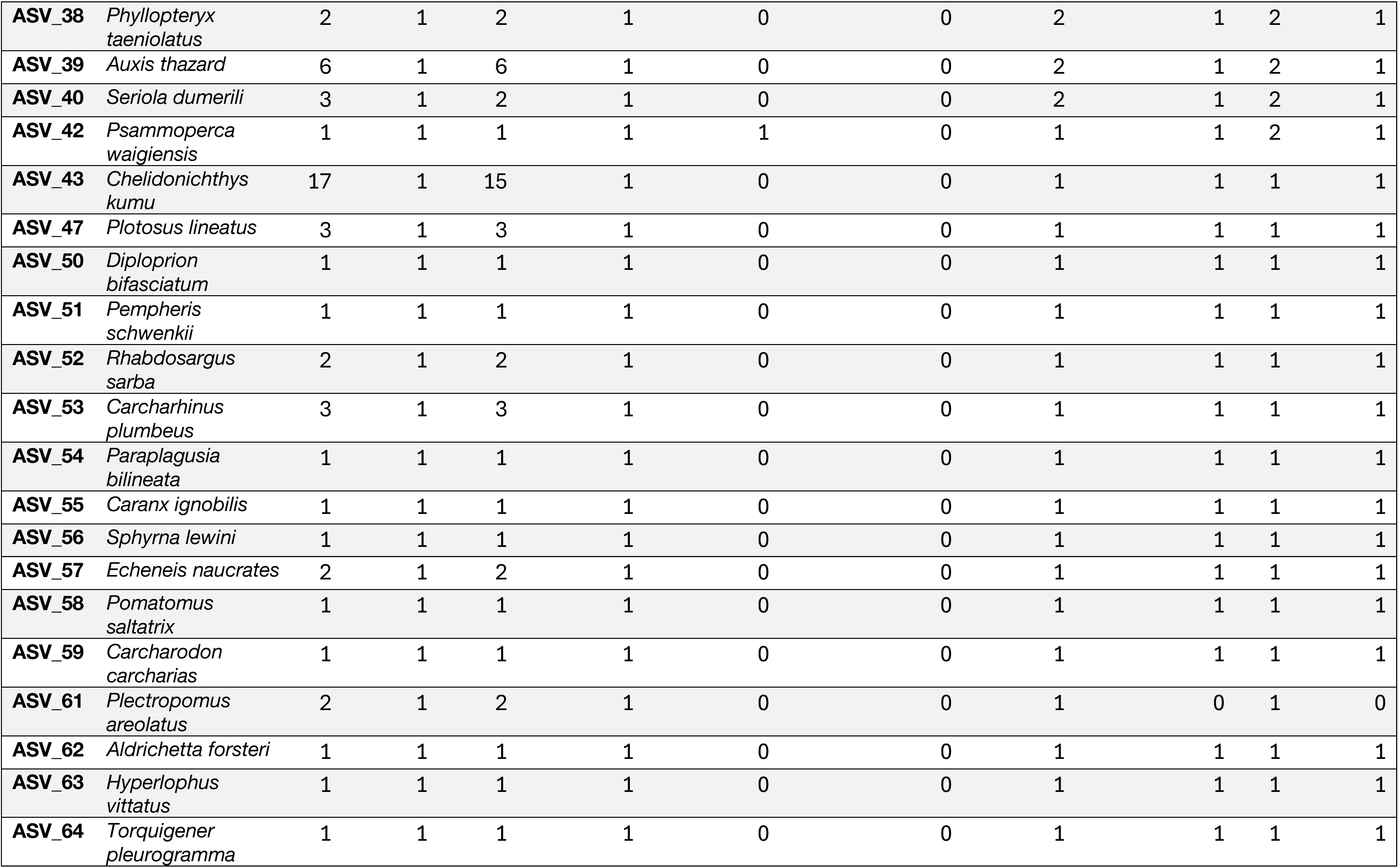
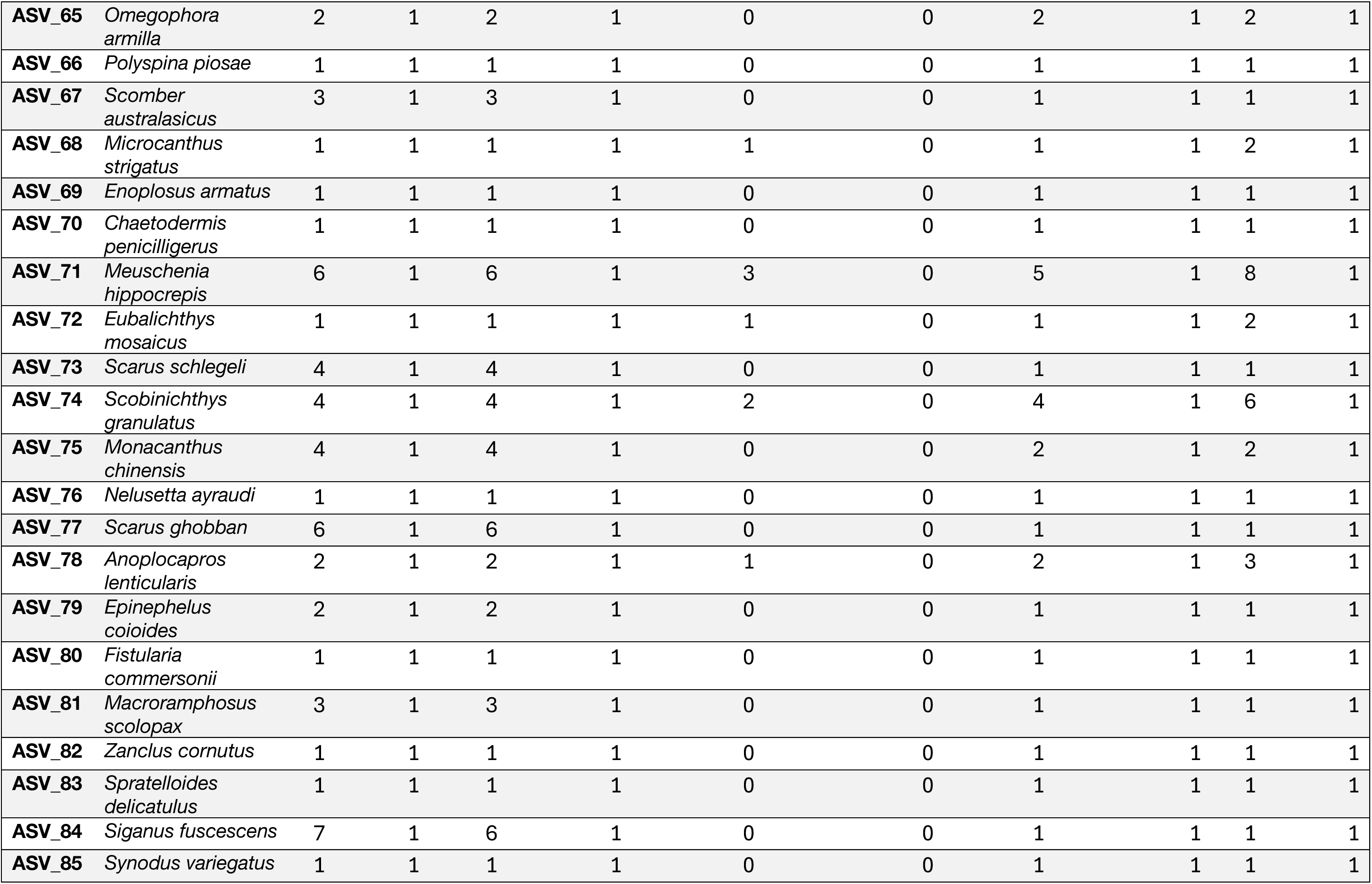
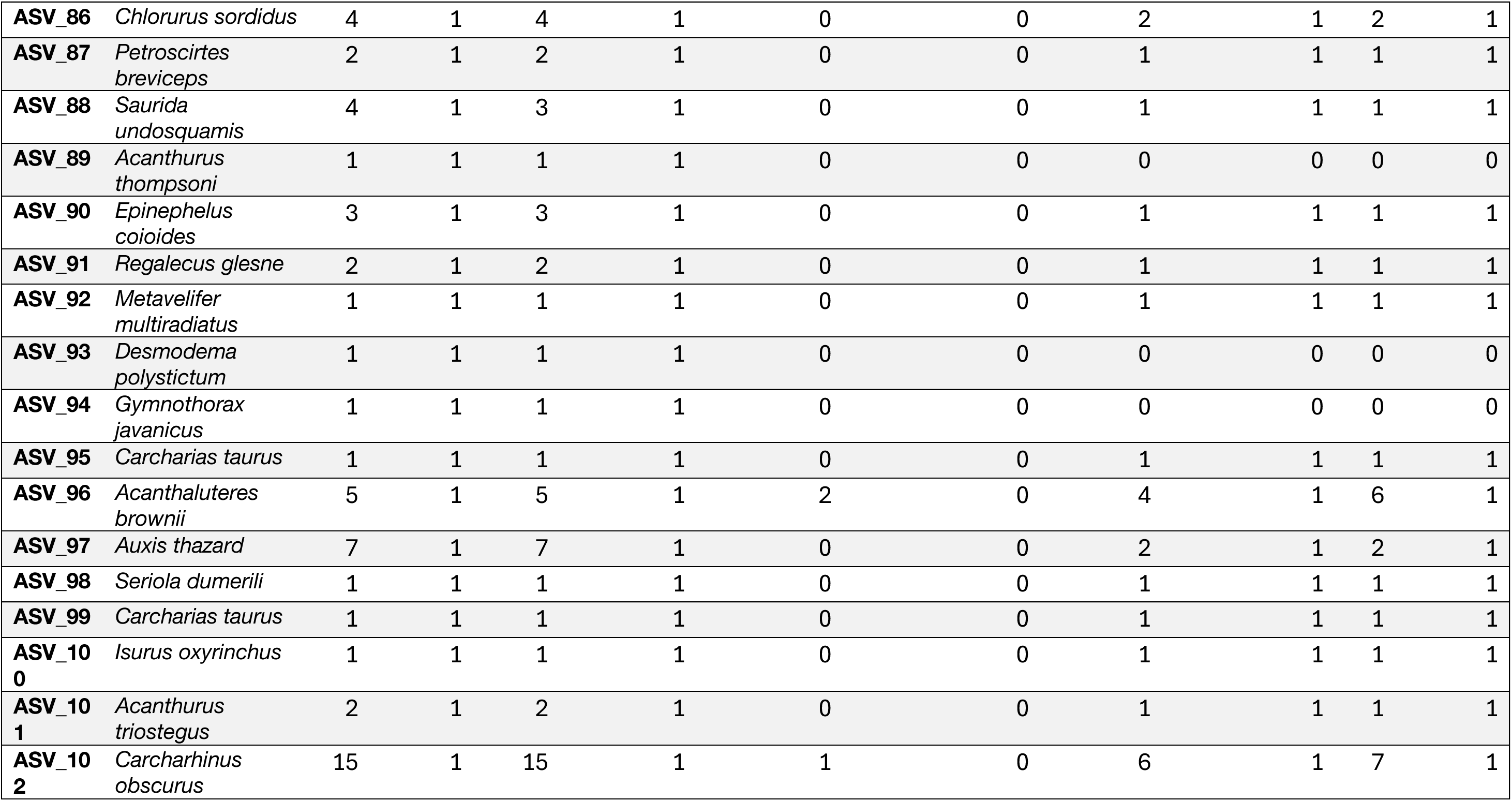

| **Sequence_id** | **TRUTH** | **MOL** | **MOL_correct** | **MOL+TAX** | **MOL+TAX_correct** | **Added by TAX+GEO** | **Added by TAX+GEO_correct** | **MOL+TAX+GEO** | **MOL+TAX+GEO_correct** | **FINAL** | **FINAL_correct** |
|---|---|---|---|---|---|---|---|---|---|---|---|
| **ASV_1** | *Phyllopteryx taeniolatus* | 1 | 1 | 1 | 1 | 0 | 0 | 1 | 1 | 1 | 1 |
| **ASV_2** | *Phycodurus eques* | 1 | 1 | 1 | 1 | 0 | 0 | 1 | 1 | 1 | 1 |
| **ASV_3** | *Phyllopteryx taeniolatus* | 1 | 1 | 1 | 1 | 0 | 0 | 1 | 1 | 1 | 1 |
| **ASV_5** | *Isurus oxyrinchus* | 1 | 1 | 1 | 1 | 0 | 0 | 1 | 1 | 1 | 1 |
| **ASV_6** | *Seriola lalandi* | 4 | 1 | 3 | 1 | 0 | 0 | 1 | 1 | 1 | 1 |
| **ASV_7** | *Phyllopteryx taeniolatus* | 1 | 1 | 1 | 1 | 0 | 0 | 1 | 1 | 1 | 1 |
| **ASV_8** | *Zeus faber* | 1 | 1 | 1 | 1 | 0 | 0 | 1 | 1 | 1 | 1 |
| **ASV_9** | *Gymnothorax pseudothyrsoideus* | 1 | 1 | 1 | 1 | 0 | 0 | 1 | 1 | 1 | 1 |
| **ASV_10** | *Carcharhinus falciformis* | 1 | 1 | 1 | 1 | 0 | 0 | 1 | 1 | 1 | 1 |
| **ASV_11** | *Pseudocaranx dentex* | 3 | 1 | 3 | 1 | 5 | 0 | 2 | 1 | 2 | 1 |
| **ASV_12** | *Carcharhinus brachyurus* | 10 | 1 | 10 | 1 | 0 | 0 | 6 | 1 | 6 | 1 |
| **ASV_13** | *Parupeneus chrysopleuron* | 1 | 1 | 1 | 1 | 0 | 0 | 1 | 1 | 1 | 1 |
| **ASV_14** | *Sphyrna lewini* | 2 | 1 | 2 | 1 | 0 | 0 | 1 | 1 | 1 | 1 |
| **ASV_15** | *Parupeneus heptacantha* | 1 | 1 | 1 | 1 | 0 | 0 | 1 | 1 | 1 | 1 |
| **ASV_16** | *Chaetodon auriga* | 8 | 1 | 8 | 1 | 0 | 0 | 2 | 1 | 2 | 1 |
| **ASV_17** | *Dascyllus trimaculatus* | 5 | 1 | 5 | 1 | 0 | 0 | 2 | 1 | 2 | 1 |
| **ASV_18** | *Gymnothorax eurostus* | 2 | 1 | 2 | 1 | 0 | 0 | 1 | 1 | 1 | 1 |
| **ASV_19** | *Acanthocybium solandri* | 2 | 1 | 2 | 1 | 0 | 0 | 2 | 1 | 2 | 1 |
| **ASV_20** | *Upeneus tragula* | 8 | 1 | 7 | 1 | 0 | 0 | 1 | 1 | 2 | 1 |
| **ASV_21** | *Hemigymnus melapterus* | 2 | 1 | 2 | 1 | 0 | 0 | 2 | 1 | 2 | 1 |
| **ASV_22** | *Chaetodon speculum* | 2 | 1 | 2 | 1 | 0 | 0 | 1 | 1 | 1 | 1 |
| **ASV_23** | *Sardinops sagax* | 2 | 1 | 1 | 1 | 0 | 0 | 1 | 1 | 1 | 1 |
| **ASV_24** | *Bathytoshia brevicaudata* | 1 | 1 | 1 | 1 | 0 | 0 | 1 | 1 | 1 | 1 |
| **ASV_25** | *Pterois volitans* | 3 | 1 | 3 | 1 | 0 | 0 | 1 | 1 | 1 | 1 |
| **ASV_26** | *Tilodon sexfasciatus* | 2 | 1 | 2 | 1 | 0 | 0 | 1 | 1 | 1 | 1 |
| **ASV_27** | *Galeocerdo cuvier* | 1 | 1 | 1 | 1 | 0 | 0 | 1 | 1 | 1 | 1 |
| **ASV_28** | *Lagocephalus sceleratus* | 1 | 1 | 1 | 1 | 0 | 0 | 1 | 1 | 1 | 1 |
| **ASV_29** | *Arothron hispidus* | 3 | 1 | 3 | 1 | 0 | 0 | 1 | 1 | 1 | 1 |
| **ASV_30** | *Tetrapturus angustirostris* | 12 | 1 | 11 | 1 | 0 | 0 | 6 | 1 | 6 | 1 |
| **ASV_31** | *Ablennes hians* | 1 | 1 | 1 | 1 | 0 | 0 | 1 | 1 | 1 | 1 |
| **ASV_32** | *Solenostomus cyanopterus* | 1 | 1 | 1 | 1 | 0 | 0 | 1 | 1 | 1 | 1 |
| **ASV_33** | *Abudefduf vaigiensis* | 10 | 1 | 9 | 1 | 0 | 0 | 3 | 1 | 3 | 1 |
| **ASV_34** | *Mugil cephalus* | 1 | 1 | 1 | 1 | 0 | 0 | 1 | 1 | 1 | 1 |
| **ASV_35** | *Abudefduf bengalensis* | 11 | 1 | 10 | 1 | 0 | 0 | 3 | 1 | 3 | 1 |
| **ASV_36** | *Acanthurus triostegus* | 1 | 1 | 1 | 1 | 0 | 0 | 1 | 1 | 1 | 1 |
| **ASV_37** | *Seriola lalandi* | 3 | 1 | 2 | 1 | 0 | 0 | 2 | 1 | 2 | 1 |
| **ASV_38** | *Phyllopteryx taeniolatus* | 2 | 1 | 2 | 1 | 0 | 0 | 2 | 1 | 2 | 1 |
| **ASV_39** | *Auxis thazard* | 6 | 1 | 6 | 1 | 0 | 0 | 2 | 1 | 2 | 1 |
| **ASV_40** | *Seriola dumerili* | 3 | 1 | 2 | 1 | 0 | 0 | 2 | 1 | 2 | 1 |
| **ASV_42** | *Psammoperca waigiensis* | 1 | 1 | 1 | 1 | 1 | 0 | 1 | 1 | 2 | 1 |
| **ASV_43** | *Chelidonichthys kumu* | 17 | 1 | 15 | 1 | 0 | 0 | 1 | 1 | 1 | 1 |
| **ASV_47** | *Plotosus lineatus* | 3 | 1 | 3 | 1 | 0 | 0 | 1 | 1 | 1 | 1 |
| **ASV_50** | *Diploprion bifasciatum* | 1 | 1 | 1 | 1 | 0 | 0 | 1 | 1 | 1 | 1 |
| **ASV_51** | *Pempheris schwenkii* | 1 | 1 | 1 | 1 | 0 | 0 | 1 | 1 | 1 | 1 |
| **ASV_52** | *Rhabdosargus sarba* | 2 | 1 | 2 | 1 | 0 | 0 | 1 | 1 | 1 | 1 |
| **ASV_53** | *Carcharhinus plumbeus* | 3 | 1 | 3 | 1 | 0 | 0 | 1 | 1 | 1 | 1 |
| **ASV_54** | *Paraplagusia bilineata* | 1 | 1 | 1 | 1 | 0 | 0 | 1 | 1 | 1 | 1 |
| **ASV_55** | *Caranx ignobilis* | 1 | 1 | 1 | 1 | 0 | 0 | 1 | 1 | 1 | 1 |
| **ASV_56** | *Sphyrna lewini* | 1 | 1 | 1 | 1 | 0 | 0 | 1 | 1 | 1 | 1 |
| **ASV_57** | *Echeneis naucrates* | 2 | 1 | 2 | 1 | 0 | 0 | 1 | 1 | 1 | 1 |
| **ASV_58** | *Pomatomus saltatrix* | 1 | 1 | 1 | 1 | 0 | 0 | 1 | 1 | 1 | 1 |
| **ASV_59** | *Carcharodon carcharias* | 1 | 1 | 1 | 1 | 0 | 0 | 1 | 1 | 1 | 1 |
| **ASV_61** | *Plectropomus areolatus* | 2 | 1 | 2 | 1 | 0 | 0 | 1 | 0 | 1 | 0 |
| **ASV_62** | *Aldrichetta forsteri* | 1 | 1 | 1 | 1 | 0 | 0 | 1 | 1 | 1 | 1 |
| **ASV_63** | *Hyperlophus vittatus* | 1 | 1 | 1 | 1 | 0 | 0 | 1 | 1 | 1 | 1 |
| **ASV_64** | *Torquigener pleurogramma* | 1 | 1 | 1 | 1 | 0 | 0 | 1 | 1 | 1 | 1 |
| **ASV_65** | *Omegophora armilla* | 2 | 1 | 2 | 1 | 0 | 0 | 2 | 1 | 2 | 1 |
| **ASV_66** | *Polyspina piosae* | 1 | 1 | 1 | 1 | 0 | 0 | 1 | 1 | 1 | 1 |
| **ASV_67** | *Scomber australasicus* | 3 | 1 | 3 | 1 | 0 | 0 | 1 | 1 | 1 | 1 |
| **ASV_68** | *Microcanthus strigatus* | 1 | 1 | 1 | 1 | 1 | 0 | 1 | 1 | 2 | 1 |
| **ASV_69** | *Enoplosus armatus* | 1 | 1 | 1 | 1 | 0 | 0 | 1 | 1 | 1 | 1 |
| **ASV_70** | *Chaetodermis penicilligerus* | 1 | 1 | 1 | 1 | 0 | 0 | 1 | 1 | 1 | 1 |
| **ASV_71** | *Meuschenia hippocrepis* | 6 | 1 | 6 | 1 | 3 | 0 | 5 | 1 | 8 | 1 |
| **ASV_72** | *Eubalichthys mosaicus* | 1 | 1 | 1 | 1 | 1 | 0 | 1 | 1 | 2 | 1 |
| **ASV_73** | *Scarus schlegeli* | 4 | 1 | 4 | 1 | 0 | 0 | 1 | 1 | 1 | 1 |
| **ASV_74** | *Scobinichthys granulatus* | 4 | 1 | 4 | 1 | 2 | 0 | 4 | 1 | 6 | 1 |
| **ASV_75** | *Monacanthus chinensis* | 4 | 1 | 4 | 1 | 0 | 0 | 2 | 1 | 2 | 1 |
| **ASV_76** | *Nelusetta ayraudi* | 1 | 1 | 1 | 1 | 0 | 0 | 1 | 1 | 1 | 1 |
| **ASV_77** | *Scarus ghobban* | 6 | 1 | 6 | 1 | 0 | 0 | 1 | 1 | 1 | 1 |
| **ASV_78** | *Anoplocapros lenticularis* | 2 | 1 | 2 | 1 | 1 | 0 | 2 | 1 | 3 | 1 |
| **ASV_79** | *Epinephelus coioides* | 2 | 1 | 2 | 1 | 0 | 0 | 1 | 1 | 1 | 1 |
| **ASV_80** | *Fistularia commersonii* | 1 | 1 | 1 | 1 | 0 | 0 | 1 | 1 | 1 | 1 |
| **ASV_81** | *Macroramphosus scolopax* | 3 | 1 | 3 | 1 | 0 | 0 | 1 | 1 | 1 | 1 |
| **ASV_82** | *Zanclus cornutus* | 1 | 1 | 1 | 1 | 0 | 0 | 1 | 1 | 1 | 1 |
| **ASV_83** | *Spratelloides delicatulus* | 1 | 1 | 1 | 1 | 0 | 0 | 1 | 1 | 1 | 1 |
| **ASV_84** | *Siganus fuscescens* | 7 | 1 | 6 | 1 | 0 | 0 | 1 | 1 | 1 | 1 |
| **ASV_85** | *Synodus variegatus* | 1 | 1 | 1 | 1 | 0 | 0 | 1 | 1 | 1 | 1 |
| **ASV_86** | *Chlorurus sordidus* | 4 | 1 | 4 | 1 | 0 | 0 | 2 | 1 | 2 | 1 |
| **ASV_87** | *Petroscirtes breviceps* | 2 | 1 | 2 | 1 | 0 | 0 | 1 | 1 | 1 | 1 |
| **ASV_88** | *Saurida undosquamis* | 4 | 1 | 3 | 1 | 0 | 0 | 1 | 1 | 1 | 1 |
| **ASV_89** | *Acanthurus thompsoni* | 1 | 1 | 1 | 1 | 0 | 0 | 0 | 0 | 0 | 0 |
| **ASV_90** | *Epinephelus coioides* | 3 | 1 | 3 | 1 | 0 | 0 | 1 | 1 | 1 | 1 |
| **ASV_91** | *Regalecus glesne* | 2 | 1 | 2 | 1 | 0 | 0 | 1 | 1 | 1 | 1 |
| **ASV_92** | *Metavelifer multiradiatus* | 1 | 1 | 1 | 1 | 0 | 0 | 1 | 1 | 1 | 1 |
| **ASV_93** | *Desmodema polystictum* | 1 | 1 | 1 | 1 | 0 | 0 | 0 | 0 | 0 | 0 |
| **ASV_94** | *Gymnothorax javanicus* | 1 | 1 | 1 | 1 | 0 | 0 | 0 | 0 | 0 | 0 |
| **ASV_95** | *Carcharias taurus* | 1 | 1 | 1 | 1 | 0 | 0 | 1 | 1 | 1 | 1 |
| **ASV_96** | *Acanthaluteres brownii* | 5 | 1 | 5 | 1 | 2 | 0 | 4 | 1 | 6 | 1 |
| **ASV_97** | *Auxis thazard* | 7 | 1 | 7 | 1 | 0 | 0 | 2 | 1 | 2 | 1 |
| **ASV_98** | *Seriola dumerili* | 1 | 1 | 1 | 1 | 0 | 0 | 1 | 1 | 1 | 1 |
| **ASV_99** | *Carcharias taurus* | 1 | 1 | 1 | 1 | 0 | 0 | 1 | 1 | 1 | 1 |
| **ASV_100** | *Isurus oxyrinchus* | 1 | 1 | 1 | 1 | 0 | 0 | 1 | 1 | 1 | 1 |
| **ASV_101** | *Acanthurus triostegus* | 2 | 1 | 2 | 1 | 0 | 0 | 1 | 1 | 1 | 1 |
| **ASV_102** | *Carcharhinus obscurus* | 15 | 1 | 15 | 1 | 1 | 0 | 6 | 1 | 7 | 1 |

